# Transcription factor residence time dominates over concentration in transcription activation

**DOI:** 10.1101/2020.11.26.400069

**Authors:** Achim P. Popp, Johannes Hettich, J. Christof M. Gebhardt

**Affiliations:** Institute of Biophysics, Ulm University, Albert-Einstein-Allee 11, 89081 Ulm, Germany

## Abstract

Transcription is a vital process activated by transcription factor (TF) binding. The active gene releases a burst of transcripts before turning inactive again. While the basic course of transcription is well understood, it is unclear how binding of a TF affects the frequency, duration and size of a transcriptional burst. We systematically varied the residence time and concentration of a synthetic TF and characterized the transcription of a reporter gene by combining single molecule imaging, single molecule RNA-FISH, live transcript visualisation and analysis with a novel algorithm, Burst Inference from mRNA Distributions (BIRD). For this well-defined system, we found that TF binding solely affected burst frequency and variations in TF residence time had a stronger influence than variations in concentration. This enabled us to device a model of gene transcription, in which TF binding triggers multiple successive steps before the gene transits to the active state and actual mRNA synthesis is decoupled from TF presence. We quantified all transition times of the TF and the gene, including the TF search time and the delay between TF binding and the onset of transcription. Our quantitative measurements and analysis revealed detailed kinetic insight, which may serve as basis for a bottom-up understanding of gene regulation.

## INTRODUCTION

Transcription of mRNA is subject to large fluctuations. Periods of transcriptional activity of a gene, during which several mRNA molecules are produced, are followed by periods of quiescence without transcription ^1^. This bursting behaviour has been observed for most investigated genes ^2,3^, and in various species ranging from bacteria ^4–6^ to yeast ^7,8^ and mammals ^9–12^. Although biological processes are intrinsically stochastic due to low molecule counts and thermal forces, the large fluctuations of transcriptional bursts point to a high degree of cooperativeness within the underlying molecular mechanisms. Indeed, numerous regulatory factors of transcriptional bursting have been observed, including enhancer-promoter interactions ^13,14^, DNA topology ^4,15^, chromatin modifications and chromatin remodelers ^16–18^, co-activators such as p300, CBP and mediator ^15^, assembly of general transcription factors and the preinitiation complex, polymerase pausing and reinitiation ^17,19–21^. Not least, binding of specific transcription factors (TFs) to cis regulatory sequences, usually representing the first step in the cascade of transcription activation,is associated with bursting ^2,7,11,16,17,22–31^.

A bursting gene can usually be described by the two-state or random-telegraph model ^32^. There, switching of the gene between the quiescent and the active state is modelled as stochastic process with on- and off-rates, and a transcription rate describes mRNA production from the active state. First insight into the mechanism of a regulatory trait, such as enhancer action or TF binding, can be obtained by revealing which of the rates of the two-state model it influences. For TFs, it is well known that TF concentration is coupled to transcriptional activity ^6,26,27,33,34^. Mostly, increasing concentration was associated with an increase in burst frequency ^2,16,26,27,30^. An effect on burst duration ^29^ or burst size ^2,10^ was also reported, although care has to be taken to avoid concatenating individual bursts ^35^.

Recently, in addition to concentration, TF residence time has been observed to affect bursting ^16,22,23,28,30,31^. As with concentration, longer residence time was associated with an increase in burst frequency ^16,17,28,30,31^. However, in many studies this effect was not exclusive, and an increase in burst duration ^16,17,30,31^ and sometimes in burst size ^30,31^ was also coupled to longer residence times. This mechanistic variation might trace back to the dissimilar methods of varying TF residence time. Different residence times were either achieved indirectly by studying modified TFs prevalent in certain cellular conditions ^28,30,31^, TF mutants with potentially different DNA binding mechanisms ^22^, by studying different target sequences ^16^ or by knocking out co-factors ^17^. Overall, a clear understanding of how TF binding, in particular TF residence time and concentration, affect transcriptional bursting, and which of the rates of the two-state model are altered by a TF, is still missing.

Here, we decipher the role of TF residence time and concentration on bursting of a minimal reporter gene with well-defined promoter structure. We used the Auxin-Inducible Degron (AID) system ^36^ to vary TF concentration without further genomic modification. To vary the residence time with minimal disturbances to the TF, we utilized the modular DNA binding domain of transcription-activator like effectors (TALEs) ^23^. We quantified mRNA production and bursting parameters with single molecule Fluorescence In Situ Hybridisation (smFISH) in fixed cells, live measurements of bursting with the MS2 system, and a novel analysis tool, Bursting Inference from mRNA Distributions (BIRD). We found that both TF residence time and concentration solely affected the on-rate of the gene, leaving burst size and burst duration unaffected. Interestingly, variation of TF residence time had a stronger influence than an equal variation of concentration. We could explain our observations in terms of an extended three-state model of gene transcription, in which binding of the TF switches the gene from a quiescent to a primed state, from which multiple successive transitions have to be traversed before transcription from the active state can be initiated. Overall, our measurements and analysis reveal detailed mechanistic and kinetic information on transcription of a minimal mammalian gene, which may serve as basis for a bottom-up approach in understanding regulatory traits of gene transcription.

## RESULTS

### Reporter cell lines to study TF residence time and concentration-dependent transcription

We aimed at studying the influence of TF binding on gene transcription, with a focus on the leverage of TF residence time and concentration and minimal blurring by other traits such as heterogeneous TF sites. Thus, we designed a reporter gene with well-defined promoter structure including a TATA box and BRE and Inr sites based on a minimal CMV promoter (Figure 1A and Supplementary information) ^37^. This promoter exhibits minimal background expression and high activation potential. We integrated a single copy of the Tet-operator (TetO) sequence, which is not present in the human genome, to exclude cooperative effects and binding competition with endogenous TFs. The artificial gene body consisted of 630 bp, followed by 24 copies of MS2 stem loops, which enable visualizing transcription events in living cells ^9,38^. As terminator, we chose the SV40 polyA sequence for an increased transcript level ^39^. We inserted the reporter gene into the human osteosarcoma cell line U2-OS using the FlipIn-system ^40^(Materials and Methods), as this line is well suited for single molecule fluorescence in situ hybridization (smFISH) ^23^. We validated the integration of a single FlipIn site via Southern Blot (Figure S1) and confirmed the positive FlipIn reaction to insert the reporter gene via gain of Hygromycin resistance, loss of Zeocin resistance, loss of lacZ activity and different PCR reactions on genomic DNA (Figure S2). After FlipIn of the gene construct, we stably transfected tdMCP-tdGFP for high signal-to-noise visualization of the MS2 stem loops ^41^.

**Figure 1:**
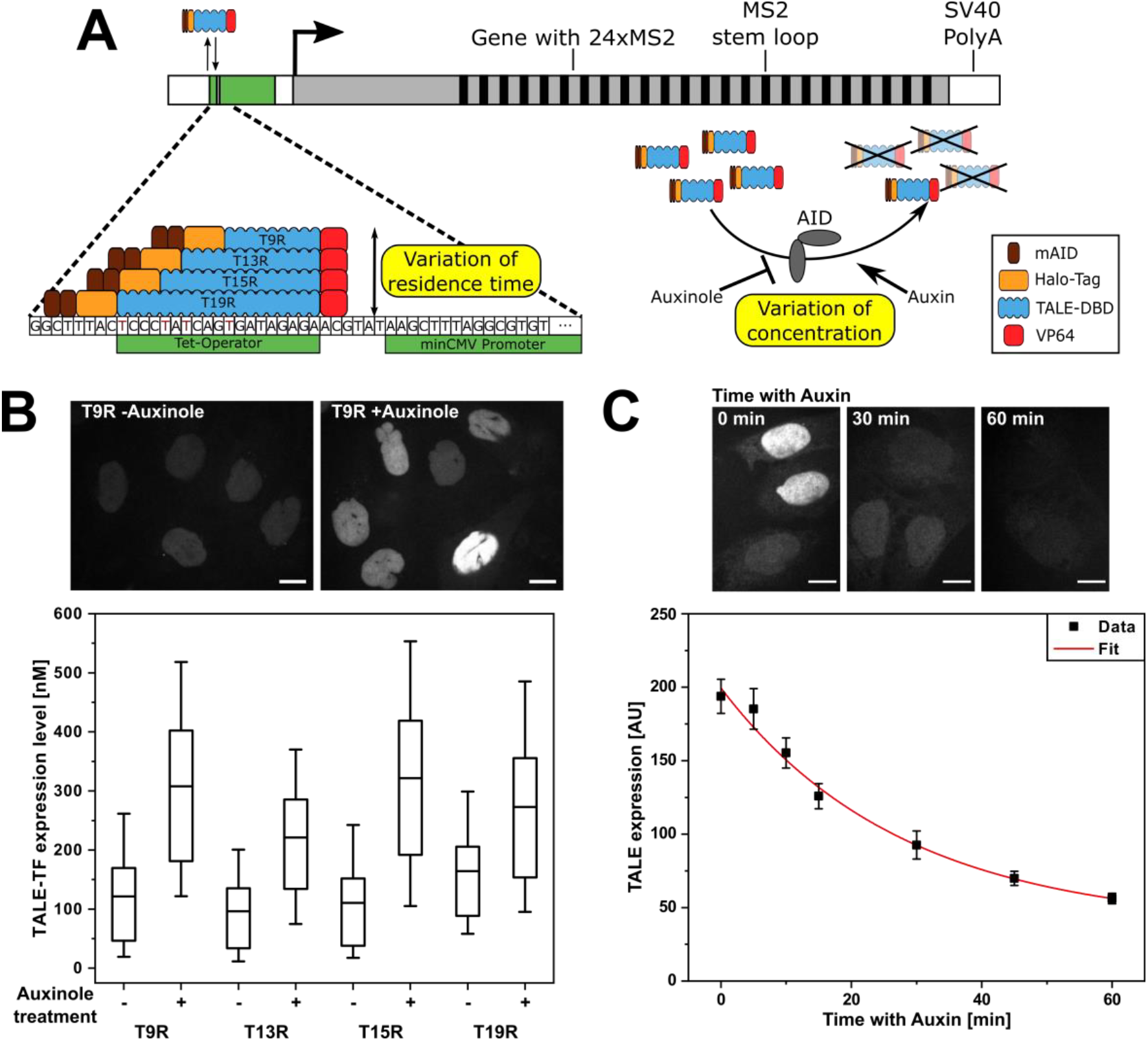
Design of reporter gene and strategy to vary TF residence time and concentration. **A)** The reporter gene consists of a TetO sequence, to which TALE-TFs are targeted, upstream of a minimal CMV-derived promoter, a 630 bp gene body, 24xMS2 repeats and a SV40 poly(A) sequence. Residence time of TALE-TFs is varied by length of the TALE DBD. Concentration of TALE-TFs is varied with the AID system and addition of either Auxinole or Auxin. **B)** Distribution of TALE-TF expression level with and without Auxinole treatment extracted from smFISH measurements. N: number of nuclei; N=450 (T9R –Auxinole); N=472 (T9R +Auxinole); N=302 (T13R –Auxinole); N=467 (T13R +Auxinole); N=476 (T15R –Auxinole); N=672 (T15R +Auxinole); N= 582 (T19R –Auxinole); N= 590 (T19R +Auxinole). Mean (line), 25^th^/75^th^ percentile (box) and 10^th^/90^th^ percentile (whiskers) define features of box plot. Inset: Fluorescence images of T9R expression without Auxinole (left) and after 16 h of Auxinole treatment (right). Scale bars denote 10 μm. **C)** TALE-TF expression level measured by spinning disc micoscopy and corrected for background as function of time after Auxin treatment (black squares) and mono-exponential fit (red line). (N: number of nuclei; N=207 (0min); N=207 (5min); N=215 (10min); N=204 (15min); N=205 (30min); N=210 (45min); N=213 (60min); N=138 (background)). Error bars denote s.e.m.. Inset: Fluorescence images of T15R expression without Auxin treatment and after 30 min and 60 min of Auxin treatment. Scale bars denote 10 μm.

As TFs, we designed artificial activators based on transcription activator like effectors (TALEs) ^42^, which allow targeting any DNA sequence starting with thymine by varying their modular DNA binding domain (TALE-DBD) ^43^. By altering the length of the TALE-DBD, TFs with different residence times can be obtained ^23^. We therefore constructed four TALE-TFs binding to 9, 13, 15 or 19 nucleotides of the TetO sequence (called T9R, T13R, T15R and T19R; Figure 1A, target sequences in Table S1), hypothesizing that they would also exhibit would also exhibit differences in their residence time. To exclude potential effects of TF position relative to the transcription start site, all TALE-DBDs ended at the same position. As activation domain, we used a C-terminal VP64 domain to achieve a high activation potential 44. The TALE-TFs further possessed an N-terminal HaloTag 45 for fluorescence imaging and a nuclear localization signal (NLS) (Figure 1B and 1C).

To adjust the concentration of TFs without further genetic modulation, we utilized two copies of the mAID-tag of the Auxin-inducible-Degron (AID) system (Figure 1A) 36. After stably transfecting the enzyme responsible for ubiquitination, OsTIR1, we created four cell lines, each stably transfected with one of the TALE-TFs and selected for comparable expression levels (Figure 1B) (Materials and Methods). By comparing the intensities of TMR labelled HaloTag-TALE-TF in the nucleus with those of a HaloTag-CTCF cell line calibrated for molecular numbers as standard 46, we determined the molar concentration of TALE-TFs within the nucleus of the four cell lines to be in the range of 100-300 nM (Figure 1B). We could increase the expression level of TALE-TFs by treating cells with 200 μM of the AID inhibitor Auxinole for 24 h (Figure 1B) 47. TALE-TFs were depleted with a half time of 28 min by treatment with 500 μM Auxin (Figure 1C).

### TALE-TFs show differences in DNA residence time

Before characterizing the binding kinetics of the TALE-TFs, we inserted additional reporter gene sequences in each of the TALE-TF cell lines by lentiviral gene transfer (Material and Methods). The average number of integrated copies in each cell line was 5 as determined by quantifying the activation profiles (Figure S3). To visualize single TALE-TFs, we labelled the HaloTag with the photostable organic SiR dye ^48^ at low labelling densities ^49^ and applied HILO microscopy ^50^ for single molecule tracking in nuclei of live cells ^23^.

To measure the dissociation rates of TALE-TFs from chromatin, we extended the time window to long observation times using time-lapse microscopy ^51^ with 50 ms frame acquisition time and frame cycle times of 0.05s, 1s and 5s (Materials and Methods). All constructs showed binding events ranging from less than a second up to several hundred seconds (Figure 2A, Supplementary Videos 1-3 and Figure S4). We first confirmed that overall TALE-TF binding interactions linearly increased with increasing concentration and did not exhibit binding saturation (Figure S5) ^52^. We then accumulated the binding times of each time-lapse condition in survival time distributions (Figure 2B and Figure S4). Next, we extracted the dissociation rate spectra of TALE-TFs using genuine rate identification (GRID) (Figure 2B, 2C and S4) ^53^. GRID yields the dissociation rate spectrum by inverse Laplace transformation of survival time distributions and enables correcting for photobleaching by global consideration of all time-lapse conditions.

**Figure 2:**
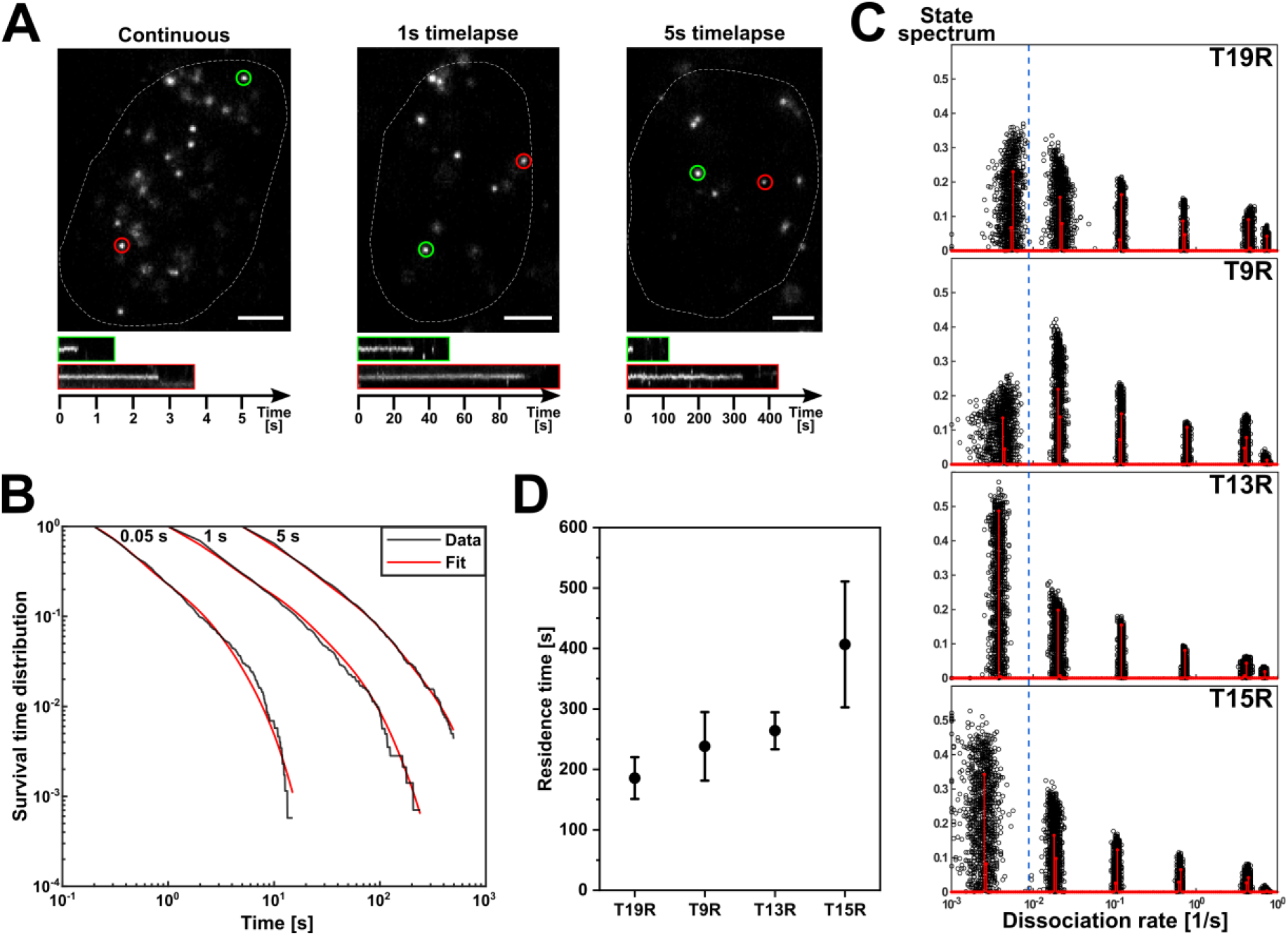
Determination of residence times of TALE-TFs. **A)** Example images of SiR-labelled HaloTag-T15R at time-lapse conditions of 0.05 s (continuous), 1 s and 5 s frame-cycle time. Images are extracted from exemplary movies S1, S2 and S3. Scale bars denote 4 μm. Lower panels: Kymographs of green and red circled molecules. **B)** Survival time distributions of SiR-HaloTag-T15R at the time-lapse condition indicated on top (black lines) and survival time functions obtained by GRID (red lines). (Number of bound molecules: 1739 (continuous); 1419 (1s time-lapse); 1608 (5s time-lapse) in 88 cells). **C)** State spectra of dissociation rates of T19R, T9R, T13R and T15R obtained by GRID using all data (red bars) and 500 resampling runs with 80% of data (black data points) as an error estimation of the spectra. **D)** Residence times of TALE-TFs extracted from the slowest dissociation rate cluster of the state spectra. Error bars denote standard deviation of the resampled spectrum.

All constructs showed 6 dissociation rate clusters and equal photobleaching rates (Figure 2C and Table S2). The 5 dissociation rate clusters corresponding to short and intermediate binding events were similar for all constructs (Figure 2C). In contrast, the dissociation rate cluster corresponding to the longest binding events differed between all constructs (Figure 2C). Since the TALE-TFs only differ by the length of their DBD, we reasoned that this dissociation rate cluster represents unbinding from the TetO sequence. We thus calculated the residence time from this dissociation rate cluster for all constructs and obtained an increasing series of residence times ranging from 186 s for T19R to 407 s for T15R (Figure 2D). We note that the variation of residence time with the length of the DBD is non-linear, similar to previous observations ^23,54^.

We further compared association of TALE-TFs to chromatin by calculating the number of binding events per area and time and normalized to concentration (pseudo-on-rate) (Figure S6) ^55^. The pseudo-on-rates of all TALE-TFs were comparable. This indicates that the length of the TALE-DBD solely modifies the longest residence time of TALE-TFs.

### TF residence time and concentration increase transcription

To quantify the number of mRNA molecules of the reporter gene in individual cells of the single reporter gene cell lines, we used smFISH (Figure 3A and Materials and Methods) ^23,56^. This approach also enables identifying sites of nascent mRNA transcription, the number of nascent mRNA molecules at those sites (burst height) and the percentage of cells exhibiting active nascent transcription (burst frequency) (Materials and Methods) ^57^. FISH probes were targeted to the SNAP gene and MS2 stem loop sequences, which are not present in the human genome and could be detected without false-positives (Figure S7). Besides mRNA, we also stained the cell membrane using Lectin-FITC and the HaloTag-TALE-TF using TMR to quantify their nuclear concentration in individual cells with U2-OS Halo-CTCF C32 ^46^ as calibrated standard (Figure 3A and Figure S8).

**Figure 3:**
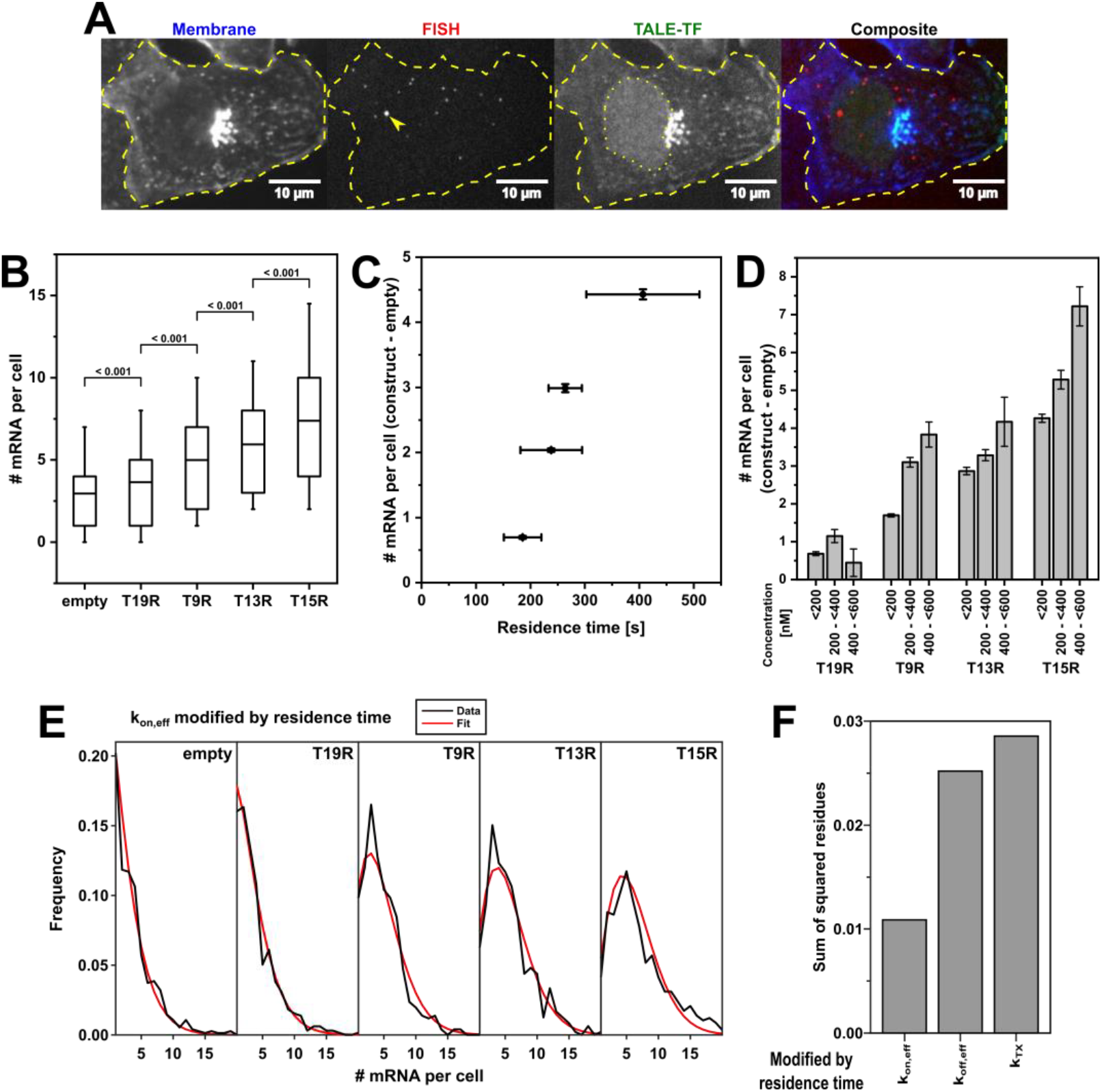
TF residence time and concentration stimulate transcription activation. **A)** Example images of the smFISH methodology: Lectin-FITC-stained cell membrane (blue; contrast 0 3000), smFISH of specific mRNA transcripts and nascent sites of transcription (arrow) (red; z-projection; contrast 2 40), TMR-stained HaloTag-TALE-TF (green; contrast 0 500) and composite image. **B)** mRNA distribution in cells expressing no TALE-TF (empty) and in cells expressing the different TALE-TF with < 300 nM. Cell numbers: N = 752 (empty); N = 888 (T19R); N = 690 (T9R); N = 664 (T13R); N = 790 (T15R). Mean (line), 25^th^/75^th^ percentile (box) and 10^th^/90^th^ percentile (whiskers) define features of box plot. Significance was determined with Wilcoxon–Mann–Whitney two-sample rank test. **C)** Mean mRNA numbers from B) of cells expressing TALE-TF subtracted with the mean mRNA number of cells expressing no TALE-TF plotted versus TALE-TF residence time. Error bars denote s.e.m. for mean mRNA numbers and resampling error estimation for residence times. **D)** Mean mRNA numbers of cells expressing TALE-TF at the indicated concentration subtracted with the mean mRNA number of cells expressing no TALE-TF. Cell numbers (<200nM, 200 - <400nM, 400 - <600nM): T19R N= 655, 375, 113; T9R N= 509, 282, 102; T13R N= 478, 256, 33; T15R N= 580, 365, 155). Error bars denote s.e.m.. **E)** mRNA distributions of TALE-TFs together with the distribution inferred by BIRD for the case where TF residence time modifies k_on,eff_ of the gene. **F)** Comparison of mRNA distributions inferred by BIRD with the measured mRNA distributions of TALE-TFs by means of the lowest sum of squared residues in cases were TF residence time modifies k_on,eff_, k_off,eff_ or k_tx_.

To determine the effect of TF residence time on transcription, we compared the distributions of mRNA molecules measured by smFISH in a cell line without TALE-TF and in the four TALE-TF cell lines (Figure 3B). In these experiments, we minimized the effect of TALE-TF concentration by only considering cells with concentrations below 300 nM. All cell lines showed different mRNA levels, with mutually significant increases from empty to T19R, T9R, T13R and T15R (Figure 3B). The mean number of mRNA produced due to TALE-TF activation above the background of leaky transcription increased with TALE-TF residence time (Figure 3C). Similar to previous findings ^16,22,23,28,30,31^, this suggests that TF residence time is a regulatory factor of transcription.

Next, we determined the effect of TF concentration on transcription. Therefore, we suppressed leaky AID-based degradation ^47^ of TALE-TFs by adding 200 μM Auxinole for 24 h before sample preparation. We excluded an effect of Auxinole on the general transcription mechanism by comparing the mRNA distributions measured by smFISH in cells without TALE-TF in absence and presence of Auxinole (Figure S9). We then quantified the mean number of mRNA molecules in each of the TALE-TF cell lines and assigned the results to 3 concentration bins (Figure 3D). As expected from previous findings ^6,26,27,33,34^, the higher the TALE-TF expression level, the higher was the number of mRNAs, again demonstrating that TF concentration is a regulatory factor of transcription.

### TF residence time and concentration solely modify the burst frequency

After having shown that both TF residence time and concentration positively correlate with transcriptionactivation, we aimed at understanding whether TF residence time affected the on-rate *kon,eff*, the transcription rate *ktx* or the off-rate *koff,gene* of the gene in a two-state model of transcriptional bursting ^6,11,32^. We therefore developed Bursting Inference from mRNA Distributions (BIRD), an inference algorithm based on iterative fitting of RNA distributions with semi-analytical solutions to the systems of differential equations describing the gene bursting kinetics (Materials and Methods). Previously, gene kinetics was simulated with the Gillespie algorithm ^58^ or modelled using hypergeometric series ^59^. We globally applied BIRD to the mRNA distributions obtained in the four TALE-TF cell lines at TF concentrations < 200 nM, while allowing only one of the rates of the two-state model to be varied at a time. Modulation of *kon,eff* by the TALE-TF residence time resulted in the best representation of our data (Figure 3E) in terms of the sum of squared residues, compared to modulating one of the other rates (Figure 3F and Figure S10).

To confirm the notion obtained from the BIRD analysis that TF residence time and concentration affect the on-rate of the reporter gene, we quantified their effect on several bursting parameters. First, we determined both frequency and height of a nascent transcription site with smFISH in each of the TALE-TF cell lines (Materials and Methods) ^57^. This approach underestimates the burst frequency and overestimates the burst height, since we only considered nascent sites with four or more transcripts to minimize false positives (Materials and Methods). Nevertheless, a relative comparison of the parameters between different conditions is possible. Consistent with the BIRD analysis, both higher TF residence time and higher concentration resulted in higher burst frequency (Figure 4A and B), while neither TF residence time nor concentration had an effect on the burst height (Figure 4A and B).

**Figure 4:**
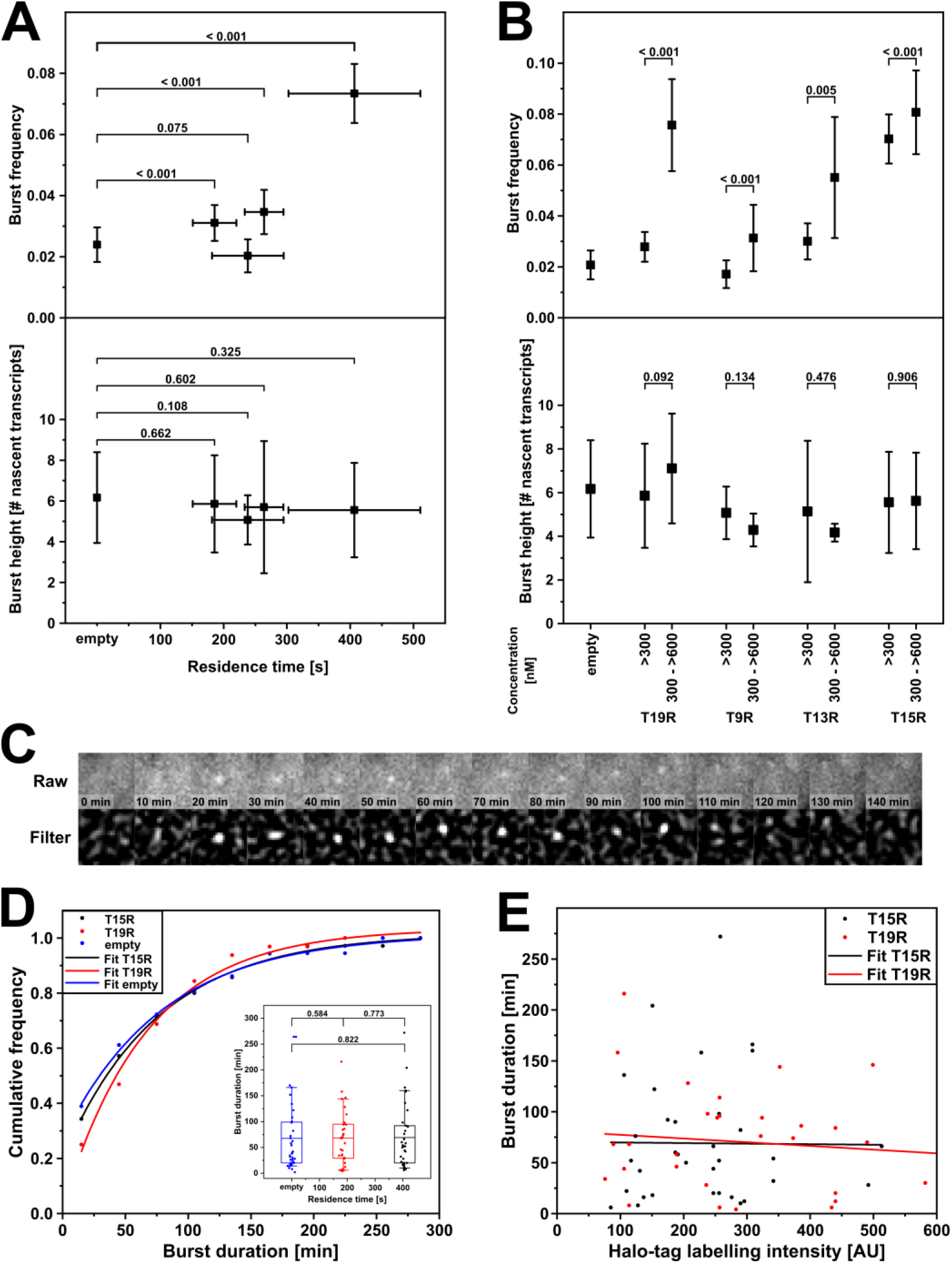
Effect of TF residence time and concentration on burst parameters. **A)** Effect of TF residence time on burst frequency and burst height. Both parameters were determined with smFISH in cells expressing < 300 nM TALE-TF. Number of detected transcription sites: N= 19 (empty); N= 28 (T19R); N= 14 (T9R); N= 23 (T13R); N= 58 (T15R). Error bars denote sqrt(N) for burst frequencies, s.d. for burst heights and resampling error estimation for residence times. Significance was tested with unranked two sample t test. **B)** Effect of TF concentration on burst frequency and burst height. Both parameters were determined with smFISH. Number of detected transcription sites (<300nM, 300 - <600nM): N= 19 (empty); N= 28, 19 (T19R); N= 14, 7 (T9R); N= 23, 6 (T13R); N= 58, 26 (T15R). Error bars denote sqrt(N) for burst frequencies and s.d. for burst heights. **C)** Example time series of a transcription site detected with the MS2 system in a living cell expressing T15R. Raw: z-projection of complete cell height; Filter: wavelet filtered z-projection. **D)** Cumulative frequency distributions of burst durations of cells expressing T15R (black symbols), T19R (red symbols), or no TALE-TF (empty, blue symbols). Lines represent mono-exponential fits. Number of transcription events: N= 35 (T15R), N= 32 (T19R), N=36 (empty). Inset: burst duration as function of TF residence time. Mean (line), 25^th^/75^th^ percentile (box) and 10^th^/90^th^ percentile (whiskers) define features of box plot. Significance was tested with Wilcoxon–Mann–Whitney two-sample rank test. **E)** Scatter plot of burst duration versus TF concentration of cells expressing T15R (black symbols) or T19R (red symbols). Lines represent linear fits. (Number of transcription events: N= 35 (T15R) and N= 32 (T19R).

The burst frequency changes if either the on-rate or the off-rate of the gene is altered, resulting in either more or longer bursts, respectively. We thus next applied the MS2 system to visualize transcription bursts in living cells and thereby directly assessed their duration (Figure 4C, Supplementary Video 4 and Materials and Methods) ^9^. Burst durations lasted for several minutes up to several hours for the TALE-TFs with longest (T15R) and shortest (T19R) residence time, with similar average burst duration of 69 min for T15R and 68 min for T19R (Figure 4D). The cumulative frequency distributions of burst durations were well described by a mono-exponential function for both T15R and T19R, indicating a single rate-determining step for the termination of bursts (Figure 4D). We also determined the effect of TF concentration on the burst duration, by quantifying the TMR-HaloTag labelling intensity of T15R and T19R (Figure 4E). As with TF residence time, burst duration was also independent of TF concentration.

We tested whether burst duration was indeed completely independent of the TF by measuring the burst duration of leaky reporter gene transcription in the cell line without TALE-TF. As predicted, the average burst duration (68 min) and the cumulative frequency distribution of burst durations were comparable to the duration and distribution in presence of TF (Figure 4D). Overall, our experiments confirm the BIRD analysis and suggest that binding of a TF solely affects the on-rate *kon,eff* of the reporter gene.

### Long TF residence time more efficiently activates transcription than frequent TF binding

Since both high TF residence time and high TF concentration increase transcription, the question arises whether it is more important to have long residence times or frequent association events of TFs for efficient activation of transcription. To answer this question, we calculated the fold-changes in RNA production for each difference in residence time of the four TALE-TF constructs, and for each difference in concentration of the different concentration bins (Figure 5A). A certain fold-change in TF residence time affects a larger fold-change in RNA production compared to the same fold-change in TF concentration. To better compare the severity of this effect, we normalized each RNA fold-change by the underlying parameter fold-change (Figure 5A) and found a 4 times larger effect for TF residence time.

**Figure 5:**
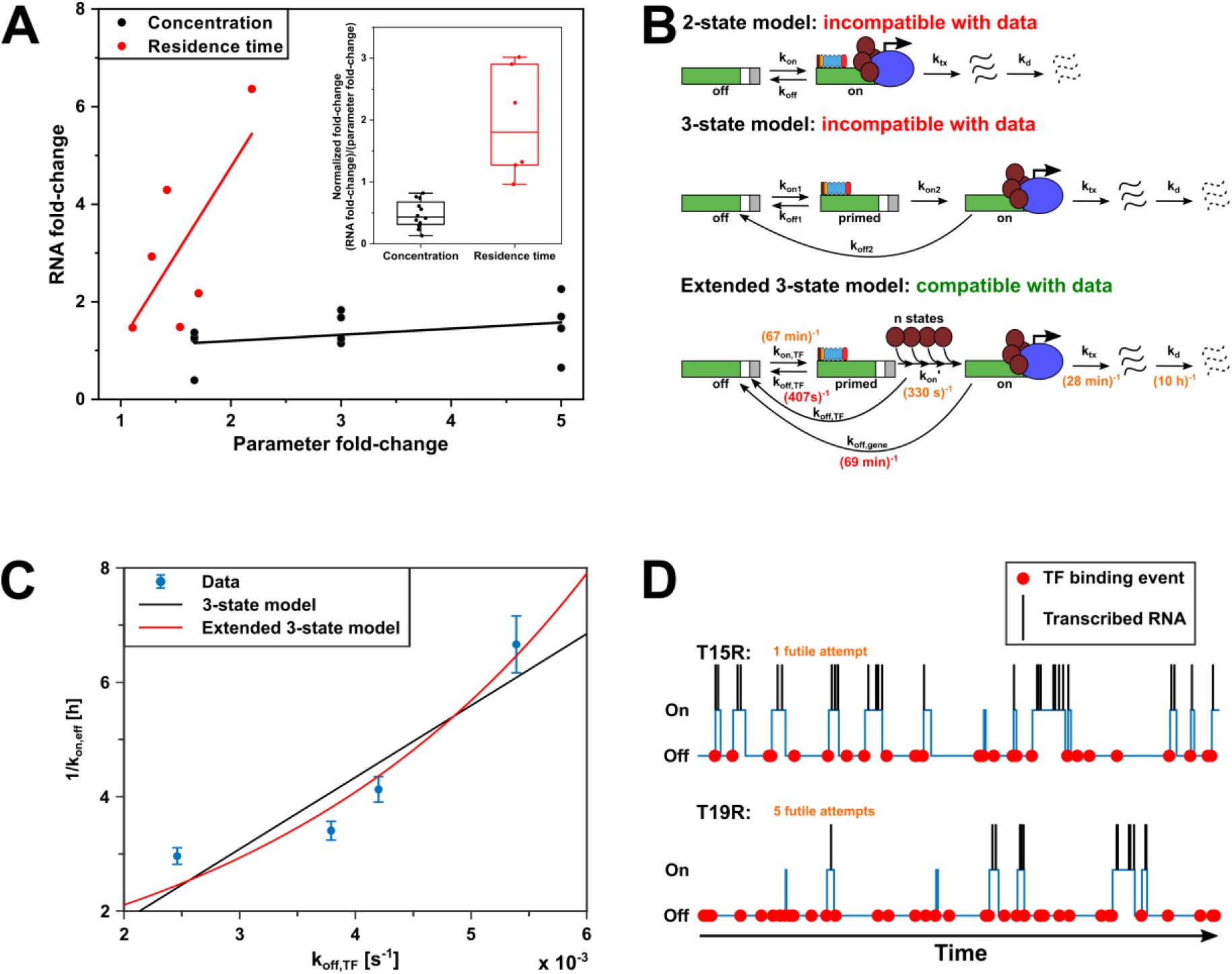
TF residence time dominates over concentration in transcription activation. **A)** Effect of TF residence time and concentration on mRNA fold-change. mRNA fold-change was calculated from mean mRNA levels (construct – empty). Residence time fold-changes were calculated for each TALE-TF combination. Concentration fold-changes were calculated from the middle values of the 3 concentration bins shown in Figure 3D. Inset: RNA fold-change normalized to corresponding parameter fold-change. **B)** Sketches of the 2-state, 3-state and extended 3-state models of gene transcription. Rates indicated in the extended 3-state model were extracted from measurements (red font) or from BIRD and modeling (orange font). **C)** Plot of 1/k_on,eff_ as obtained by BIRD versus k_off,TF_ (blue symbols). Lines represent a linear fit (black) and an exponential fit (red). Error bars denote 95% confidence interval from BIRD inference. **D)** Simulations of TF binding (red spheres), gene switching between off / primed and on-state (blue line) and transcriptional bursts (black lines) for T15R and T19R with transition rates of the extended 3-state model taken from measurements, BIRD and modelling (Supplementary Table S3). The average number of futile TF binding events before successful transcription is indicated.

The two-state model consisting of an off-state and an on-state of the gene, from which transcription occurs, well describes the RNA distribution of gene transcription (Figure 3E) ^6,11,32^. However, it is not detailed enough to include TF dissociation, in particular as the off-rate of the gene is not equal to or determined at all by binding of the TF (Figure 5B). The next simple model, a three-state model that includes a primed state from which transcription occurs upon binding of the TF and that switches to the off-state upon TF dissociation, predicts equal importance of TF residence time and TF concentration for transcription activation. To explain a dominant effect of TF residence time over TF concentration, we thus considered a multi-state model with n successive states between the primed state and the on-state of the gene (Figure 5B and Supplementary Information). In the case of large n, this model converges to the extended three-state model that predicts an exponential increase of the time necessary to switch from the off-state to the on-state with decreasing TF residence time (Supplementary Information). Indeed, the relation between 1/*kon,eff* and the dissociation rate *koff,TF* of our TALE-TFs is incompatible with a linear relation predicted by the three-state model (R^2^ = 0.84), but well described by the exponential behaviour of the extended three-state model (R^2^ = 0.94) (Figure 5C).

By combining our values for the TF residence time and the burst duration with the values for the effective on-rate, the off-rate and the transcription rate of the gene obtained by BIRD analysis of the RNA distributions, we could assign rates and values to all the transitions and parameters of the extended three-state model (Figure 5C and Supplementary Information). We found an on-rate of the TF to the gene, k_on,TF_ of (67 min)^−1^, burst frequencies of 0.12, 0.15, 0.22, 0.25 and 0.28 for empty, T19R, T9R, T13R and T15R, a transcription rate k_tx_ of (28 min)^−1^, an average burst size of 2.5 transcripts and a degradation rate k_d_ of (10 h)^−1^ (Supplementary Table S3). The values of burst frequency and burst size determined with BIRD are not blurred as those obtained by the smFISH measurement and approximate the true values. Interestingly, it takes ~330 s to pass through the n states following TF binding, which is comparable to the TF residence time. This yields mechanistic insight into the importance of the TF residence time: While the long binding T15R on average undergoes 1 futile binding event until complete transition to the on-state of the gene and successful transcription activation, the short binding T19R already undergoes 5 futile binding events (Supplementary Information). We confirmed insufficient gene activation by short TF residence time by simulating the transcription process according to the extended three-state model using all transition rates for both T15R and T19R (Figure 5D and Supplementary Information). T15R and T19R bind with equal frequency to the target site, but due to a lower number of futile attempts, T15R leads to a larger burst frequency than T19R.

## DISCUSSION

We quantified the influence of both TF residence time and concentration on the bursting parameters of a single well-defined reporter gene. We found that both regulatory traits solely affected the rate of switching on transcription, as expected from previous reports on the role of TF binding ^2,7,11,16,17,23–31^. Strikingly, we found that the regulatory effect of varying TF residence time was significantly more important than the effect of varying concentration. This enabled us to obtain mechanistic and kinetic details about the transcription activation process beyond the common two-state or a simple three-state description of gene transcription ^11,32^. Rather, we found an extended three-state model was compatible with our data, in which TF binding triggers several successive transitions of the system until the gene assumes the active state and productive transcription starts. Our model is compatible with recently identified control points of transcription regulation ^19^ and adds to previous considerations on three-state or multi-state models ^11,59–61^. In our model, for which we could determine all transition rate constants, presence of the TF increases the probability of the gene to transit from the off state over a primed state to the active state. However, not just the quantity of TF interactions matters, but their quality: the probability for a successful transition to the active state is more sensitive to the TF residence time than to its concentration.

The burst duration of 70 min and the small burst size we observed for the artificial reporter gene agree with burst parameters reported previously for endogenous mammalian genes ^11,15,24,27,28,30,31,60^. At first sight, the corresponding small transcription rate with one transcript produced every 28 minutes conflicts with the fast elongation speed of RNA polymerase II of ca. 1 kb/min ^12,15,62–64^. However, additional factors such as promoter proximal pausing of RNA polymerase II, and termination and release of mRNA influence the transcription rate. Pausing of RNA polymerase II shortly after the transcription start site is frequent and occurs with half-lives of ~2 to 30 min ^65^. For 3’ termination, a duration of a few minutes has been observed ^15,20^. Together, the burst parameters we observed for the reporter gene are consistent with the current picture of mammalian gene transcription.

One of the main tasks of a TF is to recruit cofactors necessary for transcription initiation ^66,67^. Such recruitment processes require time and would benefit from long residence times of the TF. Indeed, for Oct4, facilitated recruitment by long binding Sox2 has been observed ^33^. In general, successful cofactor recruitment becomes more probable the longer the TF is associated to the promoter ^68^. The activation domain of our TALE-TFs, VP64, directly interacts with several subunits of the mediator complex ^69^. Together, possible candidates for the transitional steps between primed and active states of the gene include recruitment of the mediator complex and subsequent assembly of general transcription factors and RNA polymerase II ^70^. Additional processes might include binding of histone writers and readers such as CBP, P300 and BRD4 ^71–73^. Of note, we did not observe any major mechanistic differences between leaky transcription in absence of the TF and TF-mediated gene activation. The same extended three-state model well described both cases. This suggests that the steps between the off state and the active state should be able to occur independent of the TF, albeit with much lower probability. For general TFs, a self-sufficient assembly of the pre-initiation complex is possible ^74^. Similarly, disassembly of such a complex, or loss of histone modifications, are possible candidates for the TF-independent rate-limiting step of burst termination that we observed. As a consequence of the additional cascade of steps in the extended three-state model, an exponential relationship between the TF residence time and the number of futile binding attempts of a TF before successful transition of the gene to the active state is predicted. In contrast to T19R, T15R, which binds twice as long, efficiently activated the gene, with only one futile attempt. This is comparable to the efficiency of Mbp1p in activating an endogenous gene in yeast ^7^ and reflects the advantage of a long residence time for successful transcription activation.

Given the exponential dependence of successful transcription activation on TF residence time, the question arises whether residence times of hundreds of seconds as for the TALE-TFs are generally necessary for successful transcription. Residence times comparable to the TALE-TFs have been observed for other factors ^31,52,75–79^. However, considerably shorter residence times were reported for many TFs ^22,28,33,49,80–82^. In addition, low-affinity binding sites with sub-optimal TF binding might be beneficial to distinguish for TFs within the same TF family ^83^. Thus, the cell needs ways to compensate for short residence times. Trivially, the cell could increase the TF concentration to achieve a high on-rate of the gene, yet this is resource-intensive. Another possibility to increase the on-rate of the gene is to increase the number of TF target sites at the promoter ^11^, which reduces the search time of the TF to the location ^78^. Enhancers additionally add a prominent regulatory level to increase the burst frequency of genes ^3,13,14^. Alternatively, the cell could increase the burst size or the burst duration once the gene is active. The burst size was found to be influenced by the nature of the activation domain ^30^ and the core promoter sequence ^3^. Similarly, increasing the number of TF binding sites at the promoter was shown to increase the number of nascent transcripts ^11,30^. Indeed, proximal promoters of many endogenous genes comprise multiple TF target sites, as do enhancer elements ^84–88^. With an increasing number of TF sites at promoter or enhancer, the propensity of TFs to form dynamic TF condensates via their low-complexity domains increases ^89^, as has been observed for various factors associated with transcription ^90–94^. The size of such condensates was found to correlate with transcription output ^95^ and GR hubs have the potential to prolong transcription bursts ^31^. Thus, condensate formation is an elegant way to locally increase the concentration of TFs and thereby compensate for the disadvantage of short TF residence times.

We can obtain an upper limit for the time that any of the TALE-TFs present in the nucleus needs to find the single specific target site within the promoter of the reporter gene from the on-rate of the gene and the number of futile binding events. In our experiment, this target search time is 67 min. With the concentration of TALE-TF of ca. 100 nM in a U2-OS nucleus of approximately an ellipsoidal volume of π/6*10μm*10μm*5μm, there are ca. 16,000 TALE-TF in the nucleus. Thus, the time for one TALE-TF molecule to find the target sequence is ca. 17,900 h (744 d). A 2.5x shorter search time of ca. 7,000 h (292 d) has been observed for TetR to find a single TetO site in U2-OS cells ^78^. The difference might reflect differences in the search process between both factors. For one Sox2 molecule in embryonic stem cells, a search time to find any target sequence of 377 s was reported, and estimated to be 31 d to find a single target site ^33^, ten times shorter than for TetR. Again, different search mechanisms might account for this difference, or differences in calculating the search time from the bound fraction, residence time and number of target sites for Sox2 versus from burst frequency and concentration for TALE-TF and the direct association measurement for TetR. Search times of ca. 100 s reported for p53 ^28^ and CTCF ^96^ to find any target sequence face similar challenges as Sox2 to convert to the search time of one specific site. The search time of TALE-TF is significantly longer than the search time of ca. 0.1 h for one Lac repressor to find a single operator in *E. coli* (ca. 120 s for any TF) ^97^, and the search time of ca. 5 h for one Mbp1p molecule to find a single target site in *S. cerevisiae* (ca. 50 s for any TF) ^7^. The difference in search time between bacterium and yeast presumably predominantly reflects differences in chromatin organization and to some extend in genome size. The difference between yeast and human corresponds within a factor of 3 to the difference in genome size. Consequently, mammalian cells need to scale up the TF numbers compared to bacteria and yeast to compensate for the very long search time of one TF to find a single target site.

Our data suggest that the TALE-TF on average should be already dissociated once the gene is in the active state and transcription starts, since TF residence time is comparable to the transition time from the primed to the active state and the transcription rate is low. Thus, while the TALE-TF helps in transiting the gene to the active state, most RNA transcription initiation events of a burst will occur without a TALE-TF bound to the promoter. Consistently, we observed that transcription in absence of the TF followed the same kinetic model, and the burst duration was independent of the TF and its residence time. Our observations are also consistent with a recent finding that RNA polymerase II recruitment occurs only after burst initiation and is rather unregulated ^19^, and the finding of a delay of RNA synthesis compared to GR binding of ~3 min ^31^. In contrast, recent reports in yeast suggest that TF residence time is directly coupled to the burst duration, while TF concentration affects the on-rate of the gene ^16,17,29^. Thus, transcription in yeast apparently only occurs while the TF is bound to the promoter. This is consistent with the burst duration of a few minutes in yeast ^7,16,17,29^, which is comparable to the elongation time of the gene and compatible with a few initiation events during the residence time of the TF. In comparison, typical burst durations in mammals are much longer than the TF residence time ^11,15,24,27,31^. The different effects of TF residence time in mammals and yeast point to differences in the regulatory mechanisms of transcription activation in both species. For example, it seems that co-factor recruitment and assembly of the transcription machinery is more efficient in yeast compared to mammals. This might be due to differences between components of the yeast and mammalian pre-initiation complexes, for example TFIID or mediator ^98^. It is further interesting to speculate that the ability to uncouple TF binding to and activation of the gene from RNA polymerase II recruitment enabled higher organisms to evolve enhancer elements, to which TFs predominantly bind. Our observations for the TALE-TFs predict that also an enhancer would only shortly, in the range of a few minutes, need to closely interact with a gene promoter for successful activation. Compatible with such a scenario, it has been observed that mediator condensates only transiently come into close proximity to a site of transcription while transcription continues to go on ^91^. Further studies will be necessary to appreciate to full extend the kinetic interplay of gene activation and actual gene transcription and differences in transcription regulation between yeast and mammals.

Given the similarities in burst size, burst duration and transcription rate of the artificial reporter gene to other mammalian systems, the reporter gene with one TF target site in the proximal promoter constitutes a basis for the kinetic understanding of gene transcription. In particular, the importance of the TF residence time, the delay between TF binding and the onset of mRNA transcription, and the TF-independent transcription termination will be able to inform the kinetic behaviour of endogenous mammalian genes with more complex promoter structures.

## MATERIALS AND METHODS

### Cloning

#### Gene construct

The gene construct was synthesized by GeneArt (ThermoFisher). We integrated 24x MS2 stem loops from phage-CMV-CFP-24xMS2 (Wu et al., 2012) by EagI-HF and BglII restriction and ligation. Afterwards we exchanged the plasmid backbone to pcDNA5/FRT from pcDNA5/FRT/TO V5 ^99^ using MluI-HF and and SphI-HF digestion. The sequence of the gene construct with inserted 24xMS2 can be found in the Supplementary Information.

For lentivirial gene transfer, we transferred the gene construct containing 24xMS2 repeats into the pLenti backbone of pLenti-CMV-OsTIR1 (Section CMV-OsTIR1) with restriction by MreI-Fast and XhoI. To allow this, we inserted a MreI and a XhoI restriction site into pLenti-CMV-OsTIR1 by annealing the two primers Cloning_pLenti_fw and Cloning_pLenti_rev and digestion with BamHI-HF and ClaI.

#### tdMCP-tdGFP

To generate a tandem-GFP version of phage-ubc-nls-ha-tdMCP-gfp ^41^, we amplified the GFP enconding sequence by PCR with the primers Cloning_MCP-GFP_fw and Cloning_MCP-GFP_rev and inserted it via XbaI restriction and ligation. We checked for correct orientation with sequencing.

#### CMV-OsTIR1

We amplified OsTIR1 from pMK232 ^100^ via PCR with the primers Cloning_OsTIR1_fw and Cloning_OsTIR1_rev and inserted it into the pLenti backbone from pLenti-CMV-rtTA3 (Addgene plasmid #26429) by digestion with BstXI. We checked for correct orientation with sequencing.

#### TALE-TF backbone

We modified the pICE-Halo-VP64 plasmid from ^23^ for this study. To hinder regulation of TALE-TF expression due to binding of the TALE-TF to their own promoter region, we removed the Tet-operators on the plasmid by SacI-HF digestion and ligation. Then, we inserted two repeats of the mAID tag, which we amplified via PCR on pMK292 ^100^.We therefore used for the first repeat the primers Cloning_mAID1_fw and Cloning_mAID1_rev together with HindIII-HF and MluI-HF digestion, and for the second repeat Cloning_mAID2_fw and Cloning_mAID1_rev with only MluI-HF digestion. We checked for correct orientation with sequencing. To ensure nuclear localization with those modifications, we inserted another nuclear localization signal C-terminally to the TALE-DBDs by annealing the two primers Cloning_NLS_fw and Cloning_NLS_rev and restriction with EcoRI-HF and PacI.

#### TALE-TF Golden Gate Reaction

We assembled TALE-TF with the Golden Gate TALEN and TAL Effector Kit2.0 (Addgene kit #1000000024) ^101^ as previously described ^23^. The designed TALE-TFs are listed with their target sequences in Table S1.

### Cell culture

We performed all experiments in U2-OS based cell lines, which we cultivated at 37 °C and 5% CO_2_ in DMEM supplemented with 10% FBS, 1% Glutamax, 1% nonessential amino acids, and 1% sodium pyruvate. We supplemented all antibiotics used for selection also during normal cultivation to hinder loss of the integrated sequences.

### Generation of cell lines

#### FlipIn U2-OS

We transfected U2-OS cells with linearized pFRT/lacZeo (ThermoFisher). After selection with Zeocin, we isolated colonies resulting from single cells with cloning cylinders. We screened all clones for single integration of pFRT/lacZeo by Southern Blot (Lofstrand Labs Limited) using a FRT site-specific probe ^18^.

#### FlipIn reaction

For the FlipIn reaction of the gene construct, we seeded 1.5 million FlipIn U2-OS cells on a 10 cm dish without antibiotics. After 24 hours, we transfected the cells with 9 μg pOG44 and 1 μg pcDNA5/FRT-gene construct using lipofectamine 2000. After 72 hours, we started the selection with hygromycin. After selection, we isolated single colonies with cloning cylinders and screened them for positive FlipIn via zeocin sensitivity, lack of β-galactosidase activity and PCR (Figure S2). To test for β-galactosidase activity, we lysed the cells with 0.5% Triton X-100 and incubated the lysates 24 h with a X-Gal containing buffer. A positive FlipIn reaction prohibited formation of a blue staining. For PCR, we isolated genomic DNA as described in ^102^ and we validated the site specific integration via FlipIn reaction with specific primers (FlipIn_Test1_fw, FlipIn_Test1_rev, FlipIn_Test2_fw and FlipIn_Test2_rev)) (Figure S2).

#### tdMCP-tdGFP and OsTIR1

After FlipIn of the gene construct, we stably integrated tdMCP-tdGFP and OsTIR1 into cells using a standard lentiviral production protocol (Addgene). For lentivirus production, we transfected LentiX cells with psPAX2 (Addgene #12260), pMD2.G (Addgene #12259), pLenti-CMV-OsTIR1 and phage-tdMCP-tdGFP. Thereafter, we exposed the cells for transfection to the harvested lentivirus. We selected transfected cells with blasticidin for OsTIR1 integration and sorted them via the GFP signal for tdMCP-tdGFP expression using FACS (BD FACSAria II).

#### TALE-TF

We stably transfected the TALE-TF with linearized plasmids via puromycin selection after the integration of OsTIR1 and tdMCP-tdGFP. We screened and sorted the colonies via FACS for equal expression (BD FACSAria II) of the different TALE-TF using staining with 1.25 μM HaloTag-TMR ligand following the protocol of the manufacturer (Promega).

#### Stable transfection of additional copies of gene construct

For the single molecule tracking experiments, we generated cells with multiple copies of the gene construct. Therefore, we stable transfected the cell lines containing a single copy of the reporter gene, OsTIR1, tdMCP-tdGFP and one of the TALE-TF with additional copies of the gene construct via lentiviral gene transfer using a standard lentiviral production protocol (Addgene). We quantified the number of integrations by comparing the levels of transcription activation before and after lentiviral transfection of additional copies of the gene construct.

#### AID for TALE-TF degradation

To test TALE degradation with AID, we stained the TALE-TF with 1.25 μM Halo-TMR-ligand after manufacturer protocol (Promega) in cells grown with Auxinole (Aobious) for 24 h. Directly afterwards, we exchanged the medium to DMEM supplemented with 500 μM Auxin (I2886, Sigma) to degrade TALE-TF ^100^. We than fixed the cells at different time points after addition of Auxin. For each time point, we determined TALE-TF expression with a Spinning-Disk microscope as described in (Clauß et al., 2017). We determined the background fluorescence in cells without TALE-TF after staining with Halo-TMR-ligand.

### Single molecule imaging and residence time analysis

#### Preparations

For single molecule imaging, we growed cells on glass bottom dishes (Delta T, Bioptechs). We stained the cells with 3-6 pM Halo-SiR ligand ^48^ directly before imaging according to the protocol of the manufacturer (Promega) to obtain equal molecular densities during imaging and therefore minimize differences in tracking losses.

#### Single molecule time-lapse imaging

We performed single molecule imaging as described ^23^. To distinguish dissociation rates of TALE-TFs independently from the photobleaching rate of the SiR-dye, we performed time-lapse microscopy ^51^. For each time-lapse condition, we introduced a different dark time between frame acquisitions of 50 ms exposure, resulting in frame acquisitions each 50 ms (continuous), 1 s or 5 s (time-lapses). To minimize differences in photobleaching rate, we controlled the laser power before each measurement to 200 mW/cm² We performed imaging up to 90 min per dish at 37 °C in OptiMEM. We supplemented the medium for labelling and imaging with 200 μM Auxinole (AOB8812, Aobious) to prevent degradation of TALE-TF during imaging.

#### Tracking analysis

We analyzed the single molecule microscopy data with TrackIt ^103^ to obtain fluorescence survival time distributions of bound TALE-TF. We adjusted the tracking settings for a nearest neighbour tracking algorithm to minimize false-positive connections due to nearby binding events and to obtain an equal probability for tracking losses due to tracking errors and photobleaching for all time-lapse conditions. The resulting tracking settings were SNR=5, maximal displacement of 0.6 pixels (continuous), 1.4 pixels (1 s time-lapse) and 2.0 pixels (5 s time-lapse) to separate bound from diffusing molecules and 1 gap frame for all time-lapse conditions. We used a minimal track length of 5 for continuous movies and 2 for the time-lapse conditions to minimize the effect of falsely assigned diffusing molecules to the bound population when determining the fluorescence survival time distributions of bound TALE-TF.

#### Analysis of survival time distributions using GRID

We determined the dissociation rate spectrum of bound TALE-TF by analysing the fluorescence survival time distributions obtained from time-lapse imaging with GRID ^53^. In brief, GRID performs an inverse Laplace transformation of a fluorescence survival time distribution to reveal the underlying dissociation rate spectrum. To account for photobleaching, the survival time distributions of different time-lapse measurements for a TALE-TF were analysed globally.

### Single molecule RNA-FISH (smFISH)

One day before sample preparations started, we seeded cells on 35 mm glass bottom dishes (Ibidi). We performed smFISH with additional staining of TALE-TF with 1.25 μM Halo-TMR-ligand (Promega) and of membrane with 10 μg/ml Lectin-FITC (Sigma-Aldrich) after a modified Stellaris RNA-FISH protocol (Biosearch Technologies) as described in ^23^. In brief, we hybridized fluorescently labelled probes targeting SNAP-tag and MS2 repeats to their target mRNA for 16 h to increase the signal of the target RNAs. This was followed by a total of 5 washing steps with a final washing time of 16 h to lower the background signal. We ordered the probes as a Stellaris Custom Probe set (Biosearch Technologies) and they are listed in Table S4. We analysed the smFISH data with the Matlab toolbox FISH-quant using the membrane staining to draw the cell outlines ^57^. Transcription sites were detected as brightest spot in the nucleus with a minimum of 4 transcripts to minimize false-positive detections and with a cut-off of the 5% brightest spots. We used the Halo-TMR staining to quantify nuclear TALE-TF concentrations, with U2-OS Halo-CTCF C32 ^46^ as calibrated reference (Figure S8 and Formula S1).

To achieve a higher expression of the TALE-TF in the same cell lines, we inhibited leaky degradation via the AID system ^47^ by supplementing medium with 200 μM Auxinole (AOB8812, Aobious) 24 h before sample preparation.

### MS2 measurements

For life cell transcription measurements, we seeded cells on 2-well dishes (Ibidi). To simultaneously quantifiy TALE-TF expression levels, we stained the cells directly before the measurement with 1.25 μM Halo-TMR-ligand according to the protocol of the manufacturer (Promega). We supplemented the medium for labelling and imaging with 200 μM Auxinole (Aobious) to prevent leaky degradation of TALE-TF during imaging. We omitted Halo-tag labelling and incubation with Auxinole for the cell line containing no TALE-TF.

We performed imaging with a spinning disk microscope ^23^ equipped with an additional cultivation chamber for temperature and CO_2_ control (Pecon). We imaged the Cells were in phenolred-free DMEM at 37°C and 5% CO_2_ by taking z-stacks of the entire nuclear volume (typically 12.5 μm) with a step size of 500 nm and exposure times of 200 ms every 2 min. We analysed the duration of transcriptional bursts with TrackIt. We adjusted the SNR for each cell individually depending on the brightness of the burst and the background signal. For tracking, we used a maximal displacement of 2 μm, a minimal track length of 2 and 2 gap frames to minimize premature loss of tracks.

## ACKNOWLEDGMENTS

We thank Astrid Bellan-Koch for her help with cloning. phage-CMV-CFP-24xMS2 (Addgene plasmid #40651) and phage-ubc-nls-ha-tdMCP-gfp (Addgene plasmid #40649) were gifts from Robert Singer (Albert Einstein College of Medicine, New York, USA). pcDNA5/FRT/TO V5 (Addgene plasmid #19445) was a gift from Harm Kampinga (University of Groningen, Groningen, The Netherlands). pMK232 (CMV-OsTIR1-PURO; Addgene plasmid #72834) and pMK292 (mAID-mCherry2-NeoR; Addgene plasmid #72830) were gifts from Masato Kanemakie (National Institute of Genetics, Mishima, Japan). pLenti-CMV-rtTA3 (Addgene plasmid #26429) was a gift from Eric Campeau (University of Massachusetts Medical School, Worcester, USA). pOG44 was kindly provided by David Suter (Ecole Polytechnique Fédérale de Lausanne, Lausanne, Switzerland). Halo-SiR ligand was kindly provided by K. Johnsson (Max Planck Institute for Medical Research, Heidelberg, Germany). We thank the Core Facility FACS of Ulm University for their help with cell sorting, with special thanks to Dr. Simona Ursu and Daniela Froelich. The authors thank the Ulm University Center for Translational Imaging MoMAN for its support.

## AUTHOR CONTRIBUTIONS

J.C.M.G. conceived the study; A.P.P. and J.C.M.G. designed experimental approaches; A.P.P. designed and performed cloning and cell line generation; A.P.P. performed experiments; A.P.P. and J.H. analysed data; J.H. devised the BIRD algorithm; J.H. and J.C.M.G modelled data; A.P.P. and J.C.M.G wrote the manuscript with input from all authors.

## FUNDING

The work was funded by the European Research Council (ERC) under the European Union’s Horizon 2020 Research and Innovation Program (No. 637987 ChromArch to J.C.M.G.), the German Research Foundation (SPP 2202 GE 2631/2–1 to J.C.M.G.) and the Carl Zeiss Foundation (to A.P.P.). J.C.M.G. acknowledges support by the Collaborative Research Centre 1279 (DFG 316249678).

## CONFLICT OF INTEREST STATEMENT

The authors declare no conflict of interest.

## DATA AVAILABILTY

Data supporting the findings of this manuscript will be available from the corresponding author after publication upon reasonable request. All raw single particle tracking data, mRNA distributions, burst durations and AID degradation data will be freely available after publication.

## CODE AVAILABILITY

The BIRD algorithm is freely available. A MatLab version of BIRD is available at https://gitlab.com/GebhardtLab/BIRD.

## Notes

### Competing Interest Statement

The authors have declared no competing interest.

