## Supplementary Information for "Transcription factor residence time dominates over concentration in transcription activation"

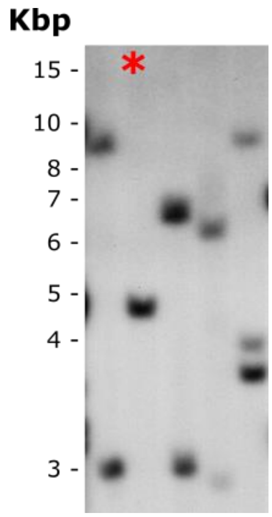


**Figure S1: Southern Blot to detect the number of integrations of pFRT/lacZeo in FlipIn U2-OS cells.** * denotes clone with single integration of FRT site taken for experiments.

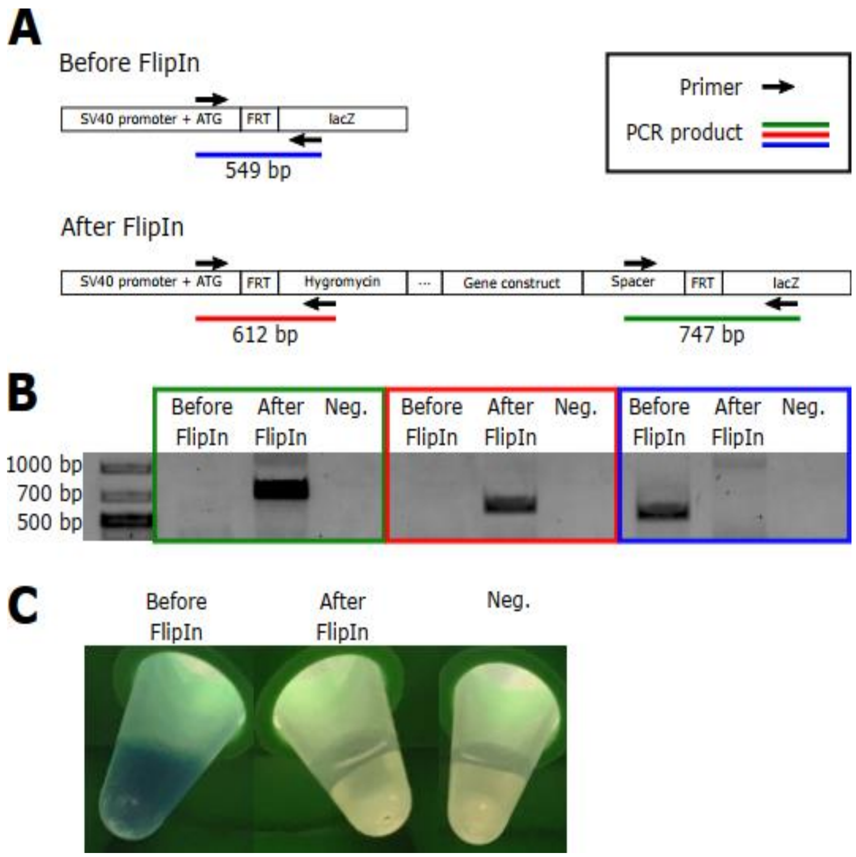


**Figure S2: Screening for positive FlipIn reaction of reporter gene into FlipIn U2-OS cells.**
A) Design of PCR tests for screening. Arrows denote primer binding sites and blue, red and green line denote PCR products.
B) PCR results of positive clone after FlipIn reaction together with negative PCR results before FlipIn reaction and negative control (no template).
C) Positive FlipIn reaction results in loss of lacZ activity and therefore no formation of blue color in X-Gal assay.

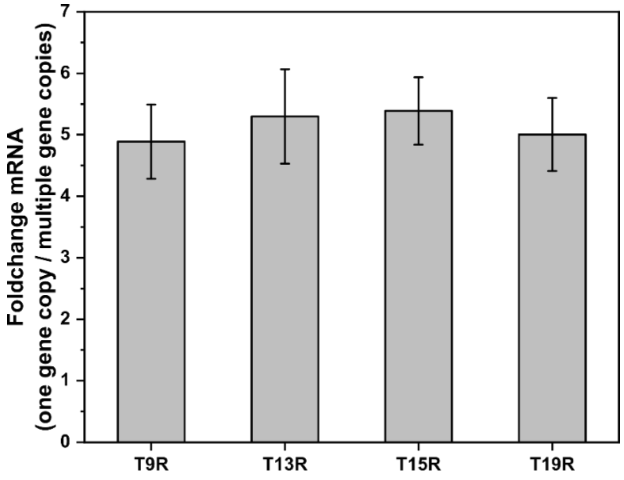


**Figure S3: Determination of the mean number of gene integrations by lentiviral gene transfer via fold change calculation of mean mRNA per cell.** Range of TALE-TF concentration <300 nM. Number of cells before lentiviral gene transfer (with one gene copy) and after lentiviral gene transfer: N= 888, 109 (T19R); N= 690, 61 (T9R); N= 664, 55 (T13R); N= 790, 110 (T15R). Error bars denote s.e.m..

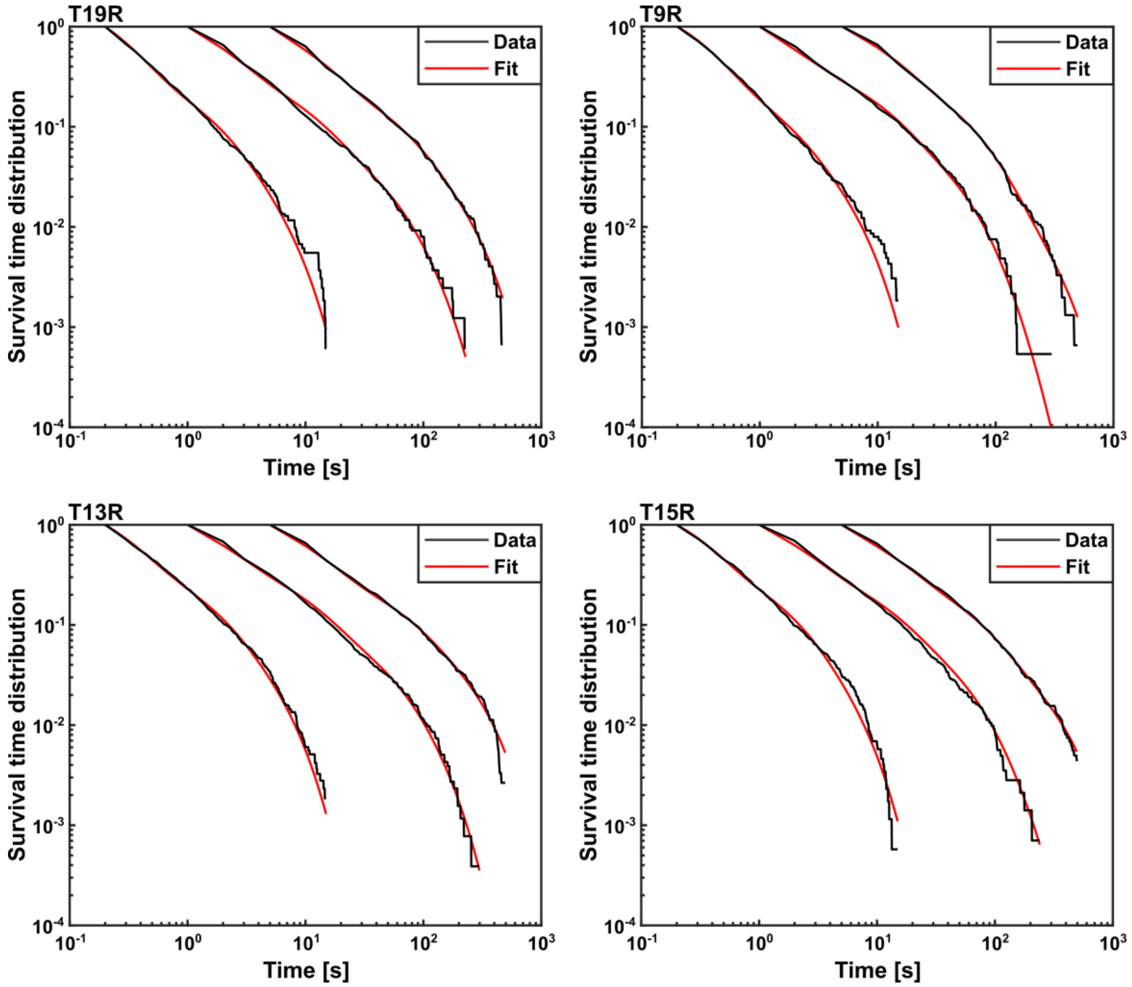


**Figure S4: Survival time distributions of T19R, T9R, T13R and T15R analyzed with GRID.** Numbers of bound molecules in continuous,1s time-lapse and 5s time-lapse, number of cells: 1632, 1626, 1486, 112 (T19R); 1631, 1858, 1519, 114 (T9R); 2152, 2570, 1507, 96 (T13R); 1739, 1419, 1608, 88 cells (T15R).

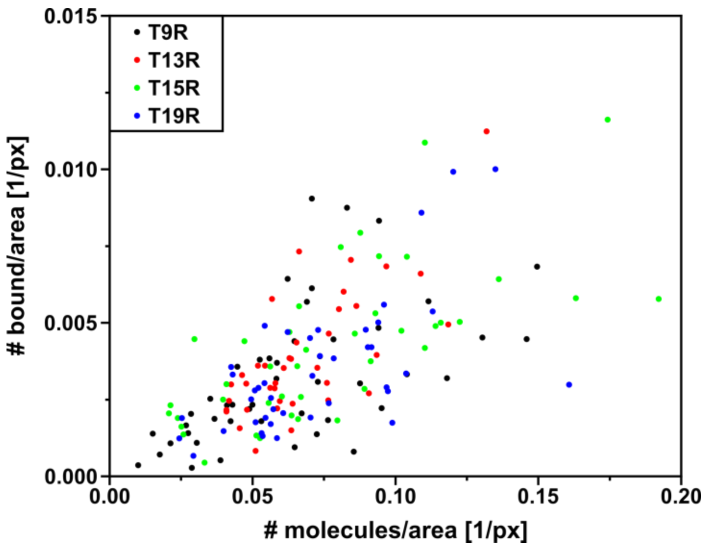


**Figure S5: Effect of TF concentration (# molecules/area) on observed binding events (# bound/area).** All data are from continuous movies. Number of molecules/cells: 33678/47 (T9R); 32320/42 (T13R); 32933/41 (T15R) 34192/42 (T19R).

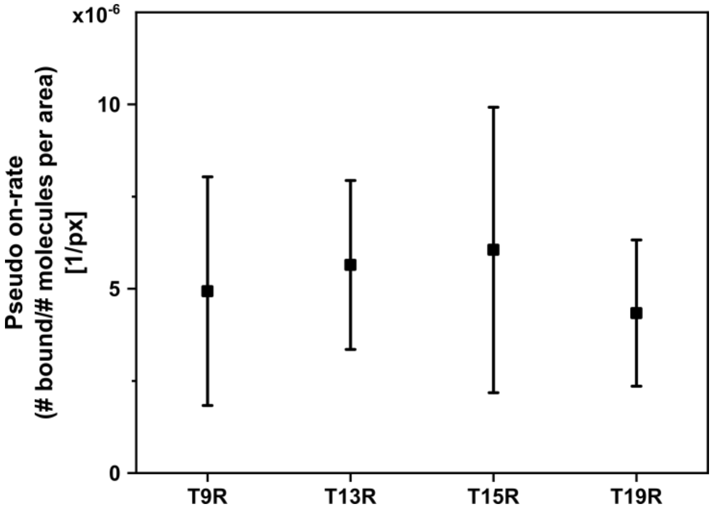


**Figure S6: Pseudo on-rate** calculated as described in (Raccaud et al., 2019). All data are from continuous movies. Number of molecules/cells: 33678/47 (T9R); 32320/42 (T13R); 32933/41 (T15R) 34192/42 (T19R). Error bars denote s.d..

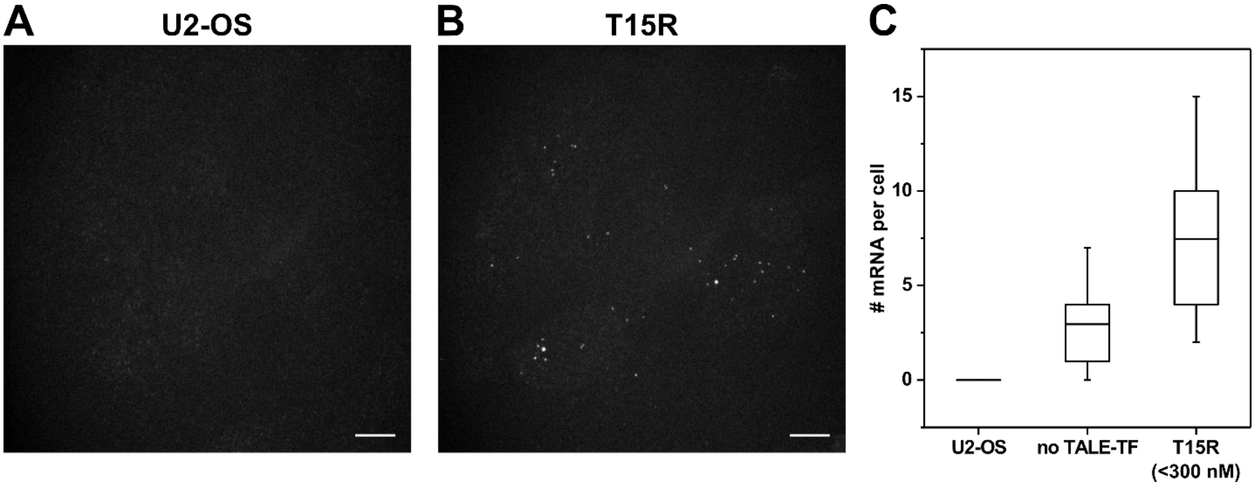


**Figure S7: smFISH in empty U2-OS cells and cells expressing the reporter gene together with T15R.**
**A)** smFISH image of U2-OS cells (z-projection; contrast 2 40). Scale bar denotes 10 µm.
**B)** smFISH image of T15R (z-projection; contrast 2 40). Scale bar denotes 10 µm.
**C)** Histograms of detected RNAs in U2-OS cells, in cells with reporter gene together with no TALE-TF and together with T15R (below 300 nM). Number of cells: N= 56 (U2-OS); N= 752 (no TALE-TF); N= 799 (T15R). Mean (line), 25$^{th}$/75$^{th}$ percentile (box) and 10$^{th}$/90$^{th}$ percentile (whiskers) define features of box plot.

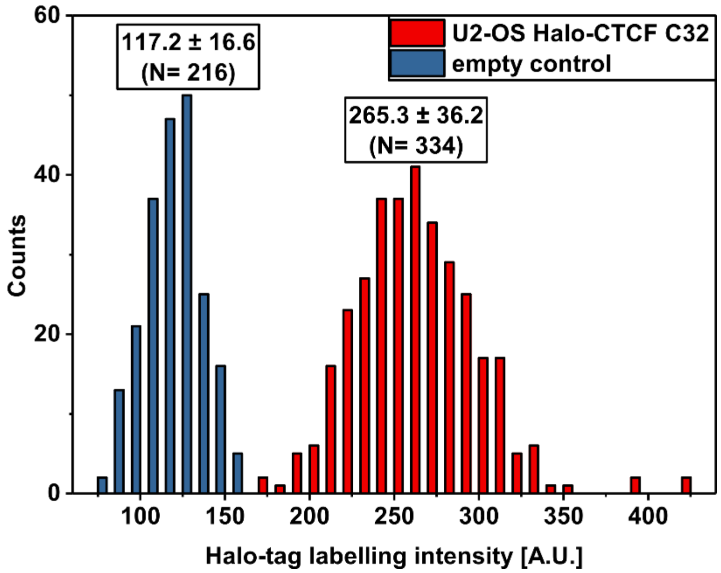


**Figure S8: Quantification of Halo-tagged proteins.** U2-OS Halo-CTCF C32 (Cattoglio et al., 2019) used as standard for quantification of Halo-tagged proteins (TALE-TF) in smFISH measurements. Nuclear Halo-TMR intensities of empty control cell line (blue) and U2-OS Halo-CTCF C32 (red) are shown. Mean intensities were taken to calculate the nuclear TALE-TF concentration with formula S1. Errors denote s.d..

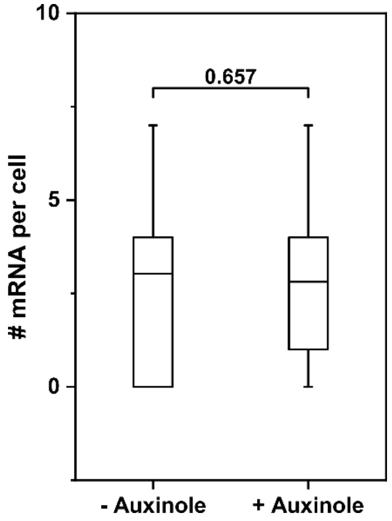


**Figure S9: Effect of Auxinole on transcription of reporter gene in absence of TALE-TF.** Significance was tested with Wilcoxon–Mann–Whitney two-sample rank test. Number of cells: N= 489 (- Auxinole) and N= 263 (+ Auxinole). Mean (line), 25$^{th}$/75$^{th}$ percentile (box) and 10$^{th}$/90$^{th}$ percentile (whiskers) define features of box plot.

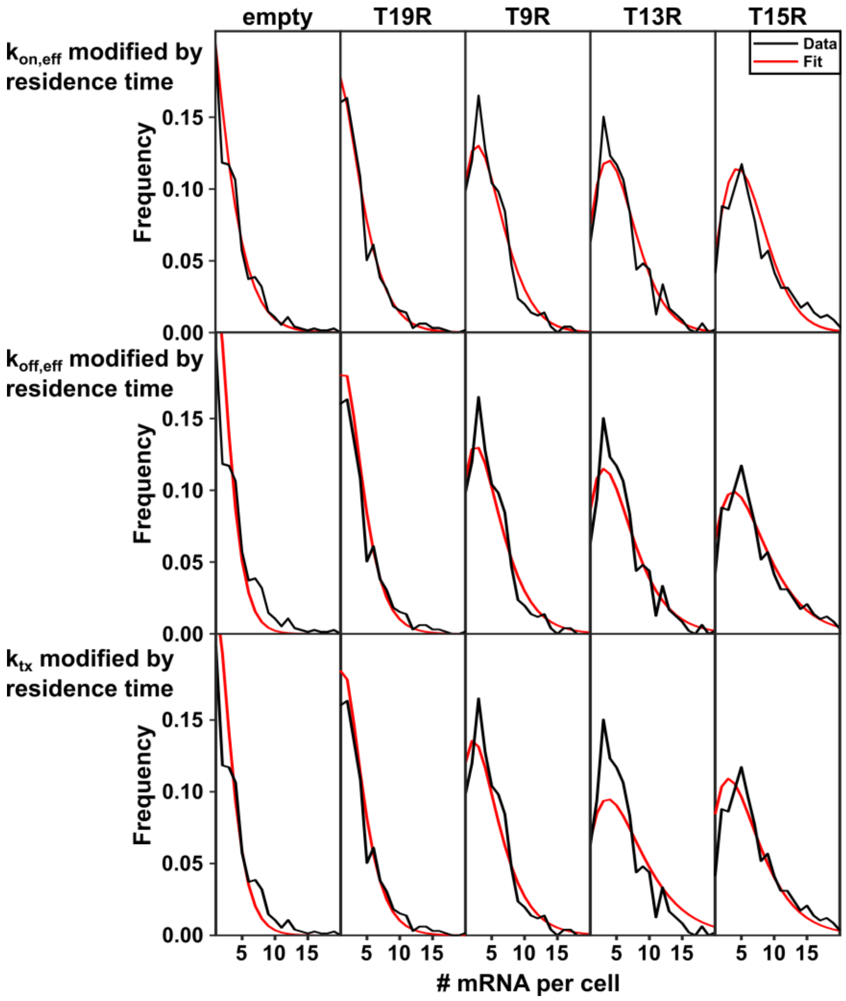


**Figure S10:** mRNA distributions of TALE-TFs (black) together with the distribution inferred by BIRD (red) for different scenarios of the effect of TF residence time.

**General analysis of RNA distributions**

Here, we develop a numerical method for fitting RNA distributions to a general multistate promoter model. In any numerical fitting procedure, the model function has to be calculated repeatedly. In case of birth and death processes, this may result in infeasible computational effort since no closed expression can be given for these systems. We therefore develop a numerical procedure for fast calculation of the RNA histograms. This procedure is then used in combination with the fmincon algorithm of the Matlab 2019b optimization toolbox to fit RNA distributions.

The state diagram for a multistate promoter model with one producing state is given by

$$\text{off 1} \underset{\mu_1}{\overset{\lambda_1}{\rightleftharpoons}} \text{off 2} \underset{\mu_2}{\overset{\lambda_2}{\rightleftharpoons}} \ldots \underset{\mu_I}{\overset{\lambda_I}{\rightleftharpoons}} \text{on} \xrightarrow{\nu} \text{RNA} \xrightarrow{\delta} \varnothing \tag{1}$$

where $\nu$ is the production rate of RNA, $\delta$ is the degradation rate of RNA, $\lambda_i, i \in 1 \ldots I$ are on-rates toward the producing state and $\mu_i$ are off-rates. Using this diagram we can give a set of differential equations for $\vec{p}(n)$,

$$\frac{d}{dt}\vec{p}(n) = \mathbf{T}\vec{p}(n) + \delta\left[(n+1)\cdot\vec{p}(n+1) - n\cdot\vec{p}(n)\right] - \nu\vec{e}_S\left[p_S(n) - p_S(n-1)\right] \tag{2}$$

where the $i$-th element of the vector is the probability to be in state $i$ while $n$ RNA molecules exist. In the above equation, we introduced the transition matrix $\mathbf{T}$ that contains the transition rates $\lambda_i, \mu_i$. The matrix element $T_{ij}$ is the rate going into state $i$ from state $j$.

We now find an iteration scheme that links $\vec{p}(n)$ with $\vec{p}(n+1)$ and thus enables us to numerically solve for $\vec{p}(n)$. First, we eliminate $p_S(n-1)$. We sum up the equations for all states and obtain

$$\frac{d}{dt}p(n) = \delta\left[(n+1)\cdot p(n+1) - n\cdot p(n)\right] - \nu\left[p_S(n) - p_S(n-1)\right] \tag{3}$$

where we introduced $p(n) = \sum_{s=1}^{S} p_s(n)$. We consider the stationary case, sum up all equations up to $n$ and obtain

$$0 = n\delta p(n) - \nu p_S(n-1) \Rightarrow \nu p_S(n-1) = n\delta p(n) \qquad \text{where} \quad n \geq 1 \tag{4}$$

Using this equation, we can eliminate $p_S(n-1)$ and obtain

$$0 = \mathbf{T}\vec{p}(n) + \delta\left[(n+1)\cdot\vec{p}(n+1) - n\cdot\vec{p}(n)\right] - \vec{e}_S\left[\nu p_S(n) - n\delta p(n)\right] \tag{5}$$

The desired iteration scheme can now be obtained by solving for $\vec{p}(n+1)$

$$\vec{p}(n+1) = -\frac{1}{\delta(n+1)}\left[\mathbf{T}\vec{p}(n) - \delta\cdot n\cdot\vec{p}(n) - \vec{e}_S\left[\nu p_S(n) - n\delta\sum_{s=1}^{S} p_s(n)\right]\right] \tag{6}$$

We define a matrix $\mathbf{M}$ such that the above equation can be rewritten as

$$\vec{p}(n+1) = \mathbf{M} \cdot \vec{p}(n) \tag{7}$$

The execution of this iteration scheme requires additional considerations since we do not know the starting values of the iteration and the iteration might diverge, depending on the eigenvalues of $\mathbf{M}$.

First, we start the iteration at the predicted expectation value $E$ of the distribution. To obtain the expectation value we calculate the probability of the gene to be in the on-state and multiply with the production rate. Doing so we avoid multiplying zeros at small and very large $n$ of left and right borders of the distribution respectively. Second, we use an eigenvector that produces eigenvalues smaller than one as our initial guess of $\vec{p}(E)$ to prevent diverging values during the iteration. After each iteration step, we renew our guess at $\vec{p}(E)$ by applying the normalization condition $\sum \sum p_s(n) = 1$.

**Fitting of measured RNA distributions**

To analyse the measured RNA distributions, we fitted a two-state model to the distributions. The model has the parameters $\lambda_{eff}, \mu_{eff}, \nu, \delta$, corresponding to $k_{on,eff}$, $k_{off,gene}$, $k_{tx}$ and $k_d$.

$$off \underset{\mu_{eff}}{\overset{\lambda_{eff}}{\rightleftharpoons}} on \xrightarrow{\nu} RNA \xrightarrow{\delta} \varnothing \tag{8}$$

The transition matrix $\mathbf{T}$ is given by

$$\mathbf{T} = \begin{pmatrix} -\lambda_{eff} & \mu_{eff} \\ \lambda_{eff} & -\mu_{eff} \end{pmatrix} \tag{9}$$

We used the gradient method fmincon of the Matlab 2019b optimization toolbox to fit the RNA histograms of all experimental conditions at once. This leads to the following problem

$$\min_{\lambda_k, \mu_k, \nu_k} \sum_{\text{Constructs}} \sum_{n=0}^{N} \left(p_n(\lambda_{eff,k}, \mu_{eff,k}, \nu_k) - \mathrm{T}_n\right)^2 \qquad k = 1\ldots5 \tag{10}$$

where the numbers 1...5 correspond to empty, T9R, T13R, T15R and T19R.

We specified boundary conditions, which correspond to different scenarios. First, we explained the histograms by an exclusive change of $\lambda_{eff}$. This corresponds to the following boundary conditions

$$\begin{aligned} \mu_{eff,k} &= \mu_{eff,l} \quad \text{where} \quad k,l = 1\ldots5 \\ \nu_k &= \nu_l \quad \text{where} \quad k,l = 1\ldots5 \end{aligned} \tag{11}$$

Second, we tried to explain the different histograms with a change in $\mu$. This corresponds to the following boundary conditions

$$\begin{aligned} \lambda_{eff,k} &= \lambda_{eff,l} \quad \text{where} \quad k,l = 1\ldots5 \\ \nu_k &= \nu_l \quad \text{where} \quad k,l = 1\ldots5 \end{aligned} \tag{12}$$

Third, we tried to explain the different histograms with a change in $\nu$. This corresponds to the following boundary conditions

$$\begin{aligned} \lambda_{eff,k} &= \lambda_{eff,l} \quad \text{where} \quad k,l = 1\ldots5 \\ \mu_{eff,k} &= \mu_{eff,l} \quad \text{where} \quad k,l = 1\ldots5 \end{aligned} \tag{13}$$

To select a model, we compared the values from equation (10) and identified the smallest value as best model.

**Derivation of the multi state and the extended 3-state models**

We discuss a multi-state model, which has an off state $p_0$ and an on-state $p_a$. These two states are separated by N intermediate states. The differential equations for this n-state model are

$$\begin{aligned} \frac{d}{dt} p_0 &= -\lambda p_0 + \mu_T \sum_{n=1}^{N} p_n + \mu p_a \\ \frac{d}{dt} p_1 &= \lambda p_0 - \mu_T p_1 - \lambda'' p_1 \\ \frac{d}{dt} p_n &= -\mu_T p_n - \lambda'' p_n + \lambda'' p_{n-1} \quad \text{where} \quad n = 2\ldots N \end{aligned} \tag{14}$$

$$\frac{d}{dt} p_a = -\mu p_a + \lambda'' p_N \tag{15}$$

where $\lambda''$ is the transition rate to higher intermediate states. All intermediate states may be depleted into the off-state with rate $\mu_T$. We assume that $\lambda'', \mu_T$ are faster than $\lambda, \mu$. Therefore, $p_n, n = 1...N$ changes faster than $p_0, p_a$ and we obtain the adiabatic approximation

$$\frac{d}{dt} p_n = 0 \quad \text{where} \quad n = 1\ldots N \tag{16}$$

As a consequence, we obtain an iteration formula for the probabilities of the intermediate states

$$\begin{aligned} p_1 &= \frac{\lambda}{\lambda''}\left(1+\frac{\mu_T}{\lambda''}\right)^{-1} p_0 \\ p_n &= \left(1+\frac{\mu_T}{\lambda''}\right)^{-1} p_{n-1} \quad \text{where} \quad n = 2\ldots N \end{aligned} \tag{17}$$

Insertion of equation (17) into equation (15) leads to

$$\frac{d}{dt} p_a = -\mu p_a + \lambda \left(1+\frac{\mu_T}{\lambda''}\right)^{-N} p_0 \tag{18}$$

We substitute the effective rate for going through all intermediate states $\lambda' = N^{-1}\lambda''$ and obtain

$$\frac{d}{dt} p_a = -\mu p_a + \lambda \left(1 + \frac{\mu_T}{N\lambda'}\right)^{-N} p_0 \tag{19}$$

The 3-state model or a single intermediate state respectively (N=1) leads to the effective on-rate

$$\begin{aligned} \frac{d}{dt} p_a &= -\mu p_a + \lambda \left(1 + \frac{\mu_T}{\lambda'}\right)^{-1} p_0 \\ \lambda_{eff} &= \lambda \left(1 + \frac{\mu_T}{\lambda'}\right)^{-1} \end{aligned} \tag{20}$$

For N ( $N >> 1$ ) intermediate states, equation (19) yields the effective on-rate of the extended 3-state model

$$\begin{aligned} \frac{d}{dt} p_a &= -\mu p_a + \lambda \left(1 + \frac{\mu_T}{N\lambda'}\right)^{-N} p_0 \\ \lambda_{eff} &= \lambda \left(1 + \frac{\mu_T}{N\lambda'}\right)^{-N} \approx \lambda \exp\left(-\frac{\mu_T}{\lambda'}\right) \end{aligned} \tag{21}$$

**Fitting of $k_{on,eff}$**

We have fitted the histograms with scenario 1 (exclusive change of $\lambda_{eff}$ ) and obtained $\lambda_{eff}$ . We plotted $1/\lambda_{eff}$ against the dissociation-rate and fitted both model (20) and (21) (3-state and N-state model derived below). We compared the residues and the R-squared values and identified the N-state model as better model.

**Number of futile binding events**

We observed that the gene on-rate ( $\lambda_{eff}$ ) gets smaller with increasing TALE-TF off-rate ( $\mu_T$ ). From this we speculated that there are futile attempts of activation if the TALE-TF is not bound long enough to enable successful transition to the active state through all intermediate states. For each TALE-TF, the number of binding events needed to successfully reach the active state can then be calculated by dividing the TALE-TF on-rate ( $\lambda$ ) by the gene on-rate ( $\lambda_{eff}$ ) (see equation (21))

$$\frac{\lambda}{\lambda_{eff}} = \exp\left(\frac{\mu_T}{\lambda'}\right) \tag{22}$$

The number of futile TALE-TF binding events is obtained by subtracting the number of binding events needed for successful activation by 1.

**Calibration of TALE-TF concentrations**

$$c_{TALE-TF} = \frac{I_{TALE-TF} - I_{empty}}{I_{C32} - I_{empty}} \times c_{C32} \quad (23)$$

with the nuclear concentration "c"; nuclear intensity of Halo-TMR "I"; 144.3 nM as $c_{32}$ (Cattoglio et al., 2019).

**Table S1. Target sequences of TALE-TF.**

| TALE-TF | Target sequence |
|---|---|
| T9R | tgatagaga |
| T13R | tcagtgatagaga |
| T15R | tatcagtgatagaga |
| T19R | tccctatcagtgatagaga |

**Table S2. Dissociation rate clusters extracted with GRID.**

| TALE-TF | Bleaching rate [$s^{-1}$] | Cluster 1 [$s^{-1}$] (%) | Cluster 2 [$s^{-1}$] (%) | Cluster 3 [$s^{-1}$] (%) | Cluster 4 [$s^{-1}$] (%) | Cluster 5 [$s^{-1}$] (%) | Cluster 6 [$s^{-1}$] (%) |
|---|---|---|---|---|---|---|---|
| T19R | 0.0111 ± 0.0010 | 7.34 ± 0.24 (4.1 ± 1.3) | 4.45 ± 0.26 (9.8 ± 1.5) | 0.707 ± 0.018 (13.3 ± 0.7) | 0.121 ± 0.006 (19.5 ± 1.0) | 0.0219 ± 0.0032 (24.2 ± 3.9) | 0.00539 ± 0.00100 (29.0 ± 4.4) |
| T9R | 0.0116 ± 0.0011 | NaN ± NaN (1.2 ± 0.9) | 4.06 ± 0.03 (12.4 ± 1.2) | 0.776 ± 0.026 (10.8 ± 0.7) | 0.119 ± 0.005 (21.8 ± 1.0) | 0.0208 ± 0.0017 (35.3 ± 2.8) | 0.00420 ± 0.00100 (18.5 ± 3.4) |
| T13R | 0.0116 ± 0.0007 | 6.89 ± 0.21 (2.0 ± 0.6) | 4.04 ± 0.24 (4.9 ± 0.7) | 0.722 ± 0.019 (8.1 ± 0.5) | 0.121 ± 0.005 (15.4 ± 0.9) | 0.0203 ± 0.0020 (20.6 ± 2.6) | 0.00379 ± 0.00044 (49.0 ± 3.2) |
| T15R | 0.0122 ± 0.0010 | 7.22 ± 0.51 (0.4 ± 0.4) | 4.25 ± 0.18 (6.6 ± 0.8) | 0.637 ± 0.019 (9.2 ± 0.9) | 0.107 ± 0.005 (14.6 ± 1.2) | 0.0183 ± 0.0019 (25.9 ± 2.3) | 0.00248 ± 0.00063 (42.4 ± 6.3) |

**Table S3. Rates for extended 3-state model. $k_{off,TF}$ and $k_{off,gene}$ were extracted from measurements, the other rates were obtained from BIRD analysis and modelling.**

| TALE-TF | $k_{on,TF}^{-1}$ [min] | $k_{off,TF}^{-1}$ [min] | $k_{on}'^{-1}$ [min] | $k_{off,gene}^{-1}$ [min] | $k_{tx}^{-1}$ [min] | $k_d^{-1}$ [h] | $k_{on,eff}^{-1}$ [min] |
|---|---|---|---|---|---|---|---|
| T19R | 67.0 | 3.1 | 5.5 | 69.0 | 28.0 | 10.0 | 394.8 |
| T9R | 67.0 | 4.0 | 5.5 | 69.0 | 28.0 | 10.0 | 266.6 |
| T13R | 67.0 | 4.4 | 5.5 | 69.0 | 28.0 | 10.0 | 232.9 |
| T15R | 67.0 | 6.8 | 5.5 | 69.0 | 28.0 | 10.0 | 150.1 |

**Table S4. FISH probes.**

| Probe # | Sequence |
|---|---|
| 1 | ttcatttcgcagtctttgtc |
| 2 | ggaagatgatacggtgcagg |
| 3 | ctggtgaaagtaggcgttga |
| 4 | taaagctctcctgctggaac |
| 5 | aacttcaccactttcagcag |
| 6 | atcagaatgggcacgggatt |
| 7 | cacagggaactcctcgatgg |
| 8 | aggtggctgtagctgatgac |
| 9 | catgttttctagagtcgacc |
| 10 | atactgcagacatgggtgat |
| 11 | aggcaattaggtaccttagg |
| 12 | catgttttctagagtcgacc |
| 13 | atactgcagacatgggtgat |
| 14 | aggcaattaggtaccttagg |
| 15 | aggcaattaggtaccttagg |
| 16 | tcatgttttctggagtcgac |
| 17 | atactgcagacatgggtgat |
| 18 | aggcaattaggtaccttagg |
| 20 | catgttttctagagtcgacc |
| 21 | atactgcagacatgggtgat |
| 22 | aggcaattaggtaccttagg |
| 24 | catgttttctagagtcgacc |
| 25 | atactgcagacatgggtgat |
| 26 | aggcaattaggtaccttagg |
| 27 | tcatgttttctggagtcgac |
| 28 | atactgcagacatgggtgat |
| 29 | aggcaattaggtaccttagg |
| 31 | catgttttctagagtcgacc |
| 32 | atactgcagacatgggtgat |
| 33 | aggcaattaggtaccttagg |
| 34 | catgttttctagagtcgacc |
| 35 | atactgcagacatgggtgat |
| 36 | aggcaattaggtaccttagg |
| 37 | catgttttctagagtcgacc |
| 38 | atactgcagacatgggtgat |
| 39 | aggcaattaggtaccttagg |
| 40 | catgttttctagagtcgacc |
| 41 | atactgcagacatgggtgat |
| 42 | aggcaattaggtaccttagg |

| | |
|---|---|
| 43 | catgttttctagagtcgacc |
| 44 | atactgcagacatgggtgat |
| 45 | aggcaattaggtaccttagg |
| 46 | catgttttctagagtccgac |
| 47 | atactgcagacatgggtgat |
| 48 | gatcagcgggtttaagatct |

**Primer list**

| **Primer** | **Sequence** |
|---|---|
| Cloning_pLenti_fw | tagcatcgatctcgagcacagtcagatcgctcgccggcgggatcctag |
| Cloning_pLenti_rev | ctaggatcccgccggcgagcgatctgactgtgctcgagatcgatgcta |
| Cloning_MCP-GFP_fw | cggctcgtctagaatggtgagcaagg |
| Cloning_MCP-GFP_rev | tatctagaatccgccttgtacagctcgtccat |
| Cloning_OsTIR1_fw | tgctaaccagtgtggtgggccaccatgacatactttcc |
| Cloning_OsTIR1_rev | cgtaatccactgtgctggatccgatggatcctcacag |
| Cloning_mAID1_fw | atcaagcttatggcaggcgccaaggagaagag |
| Cloning_mAID1_rev | cggacgcgtgctagctttatacatcctcaaatcg |
| Cloning_mAID2_fw | attacgcgtggcgccaaggagaagag |
| Cloning_NLS_fw | taacccgaaaaagaaacgcaaagtttcttg |
| Cloning_NLS_rev | aattcaagaaactttgcgtttctttttcgggttaat |
| FlipIn_Test1_fw | cgacgatacgcccatgaaga |
| FlipIn_Test1_rev | gacgacagtatcggcctcag |
| FlipIn_Test2_fw | attagtcagcaaccaggtgtgg |
| FlipIn_Test2_rev | gtcttgcaacgtgacaccct |

**Sequence of artificial gene construct after integration with the FlipIn system**

FRT site

minCMV promoter with BRE, TATA box and Inr

Tet operator

MS2 stem loop

gaagttactattccgaagttcctattctctagaaagtataggaacttccttggccaaaaagcctgaactcaccgcgacgtctgtcgagaagtttctgatc
gaaaagttcgacagcgtctccgacctgatgcagctctcggagggcgaagaatctcgtgctttcagcttcgatgtaggagggcgtggatatgtcctgc
gggtaaatagctgcgccgatggtttctacaaagatcgttatgtttatcggcactttgcatcggccgcgctcccgattccggaagtgcttgacattgggga
attcagcgagagcctgacctattgcatctcccgccgtgcacagggtgtcacgttgcaagacctgcctgaaaccgaactgcccgctgttctgcagccg
gtcgcggaggccatggatgcgatcgctgcggccgatcttagccagacgagcgggttcggcccattcggaccgcaaggaatcggtcaatacacta
catggcgtgatttcatatgcgcgattgctgatccccatgtgtatcactggcaaactgtgatggacgacaccgtcagtgcgtccgtcgcgcaggctctcg
atgagctgatgctttgggccgaggactgccccgaagtccggcacctcgtgcacgcggatttcggctccaacaatgtcctgacggacaatggccgca
taacagcggtcattgactggagcgaggcgatgttcggggattcccaatacgaggtcgccaacatcttcttctggaggccgtggttggcttgtatggag
cagcagacgcgctacttcgagcggaggcatccggagcttgcaggatcgccgcggctccgggcgtatatgctccgcattggtcttgaccaactctatc
agagcttggttgacggcaatttcgatgatgcagcttgggcgcagggtcgatgcgacgcaatcgtccgatccggagccgggactgtcgggcgtacac
aaatcgcccgcagaagcgcggccgtctggaccgatggctgtgtagaagtactcgccgatagtggaaaccgacgccccagcactcgtccgaggg
caaaggaatagcacgtactacgagatttcgattccaccgccgccttctatgaaaggttgggcttcggaatcgttttccgggacgccggctggatgatc
ctccagcgcggggatctcatgctggagttcttcgcccaccccaacttgtttattgcagcttataatggttacaaataaagcaatagcatcacaaatttca
caaataaagcatttttttcactgcattctagttgtggtttgtccaaactcatcaatgtatcttatcatgtctgtataccgtcgacctctagctagagcttggcgt
aatcatggtcatagctgtttcctgtgtgaaattgttatccgctcacaattccacacaacatacgagccggaagcataaagtgtaaagcctggggtgcct
aatgagtgagctaactcacattaattgcgttgcgctcactgcccgctttccagtcgggaaacctgtcgtgccagctgcattaatgaatcggccaacgc
gcggggagaggcggtttgcgtattgggcgctcttccgcttcctcgctcactgactcgctgcgctcggtcgttcggctgcggcgagcggtatcagctcac
tcaaaggcggtaatacggttatccacagaatcaggggataacgcaggaaagaacatgtgagcaaaaggccagcaaaaggccaggaaccgta
aaaaggccgcgttgctggcgtttttccataggctccgcccccctgacgagcatcacaaaaatcgacgctcaagtcagaggtggcgaaacccgaca
ggactataaagataccaggcgtttccccctggaagctccctcgtgcgctctcctgttccgaccctgccgcttaccggatacctgtccgcctttctcccttc
gggaagcgtggcgctttctcatagctcacgctgtaggtatctcagttcggtgtaggtcgttcgctccaagctgggctgtgtgcacgaaccccccgttca
gcccgaccgctgcgccttatccggtaactatcgtcttgagtccaacccggtaagacacgacttatcgccactggcagcagccactggtaacaggatt
agcagagcgaggtatgtaggcggtgctacagagttcttgaagtggtggcctaactacggctacactagaagaacagtatttggtatctgcgctctgct
gaagccagttaccttcggaaaaagagttggtagctcttgatccggcaaacaaaccaccgctggtagcggtggtttttttgtttgcaagcagcagattac
gcgcagaaaaaaaggatctcaagaagatcctttgatcttttctacggggtctgacgctcagtggaacgaaaactcacgttaagggattttggtcatga
gattatcaaaaaggatcttcacctagatccttttaaattaaaaatgaagttttaaatcaatctaaagtatatatgagtaaacttggtctgacagttaccaat
gcttaatcagtgaggcacctatctcagcgatctgtctatttcgttcatccatagttgcctgactccccgtcgtgtagataactacgatacgggagggctta
ccatctggccccagtgctgcaatgataccgcgagacccacgctcaccggctccagatttatcagcaataaaccagccagccggaagggccgagc
gcagaagtggtcctgcaactttatccgcctccatccagtctattaattgttgccgggaagctagagtaagtagttcgccagttaatagtttgcgcaacgtt
gttgccattgctacaggcatcgtggtgtcacgctcgtcgtttggtatggcttcattcagctccggttcccaacgatcaaggcgagttacatgatcccccat
gttgtgcaaaaaagcggttagctccttcggtcctccgatcgttgtcagaagtaagttggccgcagtgttatcactcatggttatggcagcactgcataatt
ctcttactgtcatgccatccgtaagatgcttttctgtgactggtgagtactcaaccaagtcattctgagaatagtgtatgcggcgaccgagttgctcttgcc
cggcgtcaatacgggataataccgcgccacatagcagaactttaaaagtgctcatcattggaaaacgttcttcggggcgaaaactctcaaggatctt
accgctgttgagatccagttcgatgtaacccactcgtgcacccaactgatcttcagcatcttttactttcaccagcgtttctgggtgagcaaaaacagga
aggcaaaatgccgcaaaaaagggaataagggcgacacggaaatgttgaatactcatactcttcctttttcaatattattgaagcatttatcagggttatt
gtctcatgagcggatacatatttgaatgtatttagaaaaataaacaaataggggttccgcgcacatttccccgaaaagtgccacctgacgtcgacgg
atcgggagatctcccgatcccctatggtgcactctcagtacaatctgctctgatgccgcatagttaagccagtatctgctccctgcttgtgtgttggaggtc
gctgagtagtgcgcgagcaaaatttaagctacaacaaggcaaggcttgaccgacaattgcatgaagaatctgcttagggttaggcgttttgcgctgct
tcgcgatgtacgggccagatatacgcgtcacgagactagcctcgagagttggctttactccctatcagtgatagagaacgtataagctttaggcgtgt
acggtgggcgcctataaaagcagagctcgtttagtgaaccgtcagatcgcctggagcaattccacaacacttttgtcttataccaactttccgtaccac
ttcctaccctcgtaaaaccggtcgtcgactagtccagtgtggtggaattctgcagatatcaatggacaaagactgcgaaatgaagcgcaccaccctg
gatagccctctgggcaagctggaactgtctgggtgcgaacagggcctgcaccgtatcatcttcctgggcaaaggaacatctgccgccgacgccgtg
gaagtgcctgccccagccgccgtgctgggcggaccagagccactgatgcaggccaccgcctggctcaacgcctactttcaccagcctgaggcca
tcgaggagttccctgtgccagccctgcaccacccagtgttccagcaggagagctttacccgccaggtgctgtggaaactgctgaaagtggtgaagtt

cggagaggtcatcagctacagccacctggccgccctggccggcaatcccgccgccaccgccgccgtgaaaaccgccctgagcggaaatcccgt
gcccattctgatcccctgccaccgggtggtgcagggcgacctggacgtggggggctacgagggcgggctcgccgtgaaagagtggctgctggcc
cacgagggccacagactgggcaagcctgggctgggttaaagcggccggatcctaaggtacctaattgcctagaaaacatgaggatcacccatgt
ctgcaggtcgactctagaaaacatgaggatcacccatgtctgcagtattcccgggttcattagatcctaaggtacctaattgcctagaaaacatgagg
atcacccatgtctgcaggtcgactctagaaaacatgaggatcacccatgtctgcagtattcccgggttcattagatcctaaggtacctaattgcctaga
aaacatgaggatcacccatgtctgcaggtcgactccagaaaacatgaggatcacccatgtctgcagtattcccgggttcattagatcctaaggtacct
aattgcctagaaaacatgaggatcacccatgtctgcaggtcgactctagaaaacatgaggatcacccatgtctgcagtattcccgggttcattagatc
ctaaggtacctaattgcctagaaaacatgaggatcacccatgtctgcaggtcgactctagaaaacatgaggatcacccatgtctgcagtattcccgg
gttcattagatcctaaggtacctaattgcctagaaaacatgaggatcacccatgtctgcaggtcgactccagaaaacatgaggatcacccatgtctgc
agtattcccgggttcattagatcctaaggtacctaattgcctagaaaacatgaggatcacccatgtctgcaggtcgactctagaaaacatgaggatca
cccatgtctgcagtattcccgggttcattagatcctaaggtacctaattgcctagaaaacatgaggatcacccatgtctgcaggtcgactctagaaaac
atgagaggatcacccatgtctgcagtattcccgggttcattagatcctaaggtacctaattgcctagaaaacatgaggatcacccatgtctgcaggtcg
actctagaaaacatgaggatcacccatgtctgcagtattcccgggttcattagatcctaaggtacctaattgcctagaaaacatgaggatcacccatgt
ctgcaggtcgactctagaaaacatgaggatcacccatgtctgcagtattcccgggttcattagatcctaaggtacctaattgcctagaaaacatgagg
atcacccatgtctgcaggtcgactctagaaaacatgaggatcacccatgtctgcagtattcccgggttcattagatcctaaggtacctaattgcctaga
aaacatgaggatcacccatgtctgcaggtcggactctagaaaacatgaggatcacccatgtctgcagtattcccgggttcattagatcttaaacccgc
tgatcagcctcgaaacttgtttattgcagcttataatggttacaaataaagcaatagcatcacaaatttcacaaataaagcatttttttcactgcattctagtt
gtggtttgtccaaactcatcaatgtatcttatctagagcgcgccggcggtattcagcgactcgaccaacgcaagcgtctcgcgggatttcgagcatcg
ggggttgtggtagatcgtgatcatcgttcaaggcaggaggggtgaggcctgtattgtagcgagtctgcatcgaggggcctgatgggggcaatcga
gatcggtgtggggttcctcttgataggtcgctgaggcggtttcggtttggcgatgcgcagtgatcgcgagaggggagtgtcctgtgttcggcgcgaag
agtggttgtttatgcgcataaagtcgggcgggtggcttcgcggggccgacgaccggcgcagccgatgtctcgtgagggctgtctcgtgagggctgtc
tcgtgagggctgtctcgtgagggctgtctcgggtcgcgaaaggggtatgcgattgcggaacacgccggctaggtttatgcgcataaagcgcgcgatt
gccgggcgaggtacagcgtggctcgtggcatggacatatgtcgacacgcaacggcactgcgcgatggctatatcgatagaccgtcgtccccgac
atgacgtcattgcccgcctgactcaggcgtagtctctgtcaacggcgcttcgtcgcggcctatcgttgcctctggtttatgcgcataaacactacggccc
ggcaggacattgcctccatgcacgtccattcaacgacgcacctcgtgatctacgcacctgactacgacacgcagctcacgccggacgctcgcgag
tgaataccatgcgctctagcgacgaagcccatcgatcggcctccgcagtctttatgcgcacaaactcgcggctcgcaggcaacgcggctcgtcccg
cgaaaggagagagcgtcgagatatctctgccacgaacggccccgacgatacgcccatgaagacgtaatcctgagcagggcatcgatcactcgg
catcagccgcttcctcttcaccacaccgtccaaacgagccatcgaacatccaccgctcgtttatgcgcataaaccgatcgtccgaatgggcatgctg
gggatgcggtgggctctatggcttctgaggcggaaagaaccagctggggctctaggggggtatccccacgcgccctgtagcggcgcattaagcgcg
gcgggtgtggtggttacgcgcagcgtgaccgctacacttgccagcgccctagcgcccgctcctttcgctttcttcccttcctttctcgccacgttcgccgg
ctttccccgtcaagctctaaatcgggggctccctttagggttccgatttagtgctttacggcacctcgaccccaaaaaacttgattagggtgatggttcac
gtacctagaagttcctattccgaagttcctattctctagaaagtataggaacttc
